# Succinct *k*-mer Sets Using Subset Rank Queries on the Spectral Burrows-Wheeler Transform ^*^

**DOI:** 10.1101/2022.05.19.492613

**Authors:** Jarno N. Alanko, Simon J. Puglisi, Jaakko Vuohtoniemi

**Affiliations:** University of Helsinki, Department of Computer Science, Helsinki, Finland; Dalhousie University, Faculty of Computer Science, Halifax, Canada

## Abstract

The *k*-spectrum of a string is the set of all distinct substrings of length *k* occurring in the string. This is a lossy but computationally convenient representation of the information in the string, with many applications in high-throughput bioinformatics. In this work, we define the notion of the *Spectral Burrows-Wheeler Transform* (SBWT), which is a sequence of subsets of the alphabet of the string encoding the *k*-spectrum of the string. The SBWT is a distillation of the ideas found in the BOSS and Wheeler graph data structures. We explore multiple different approaches to index the SBWT for membership queries on the underlying *k*-spectrum. We identify *subset rank queries* as the essential subproblem, and propose four succinct index structures to solve it. One of the approaches essentially leads to the known BOSS data structure, while the other three offer attractive time-space trade-offs and support simpler query algorithms that rely only on fast rank queries. The most general approach involves a novel data structure we call the *subset wavelet tree*, which we find to be of independent interest. All of the approaches are also amendable to entropy compression, which leads to good space bounds on the sizes of the data structures. Using entropy compression, we show that the SBWT can support membership queries on the *k*-spectrum of a single string in *O*(*k*) time and (*n* + *k*)(log *σ* + 1*/* ln 2) + *o*((*n* + *k*)*σ*) bits of space, where *n* is the number of distinct substrings of length *k* in the input and *σ* is the size of the alphabet. This improves from the time *O*(*k* log *σ*) achieved by the BOSS data structure. We show, via experiments on a range of genomic data sets, that the simplicity of our new indexes translates into large performance gains in practice over prior art.

## 1 Introduction

The set of substrings of a given length *k* of a string *S* is called the *k-spectrum* of *S*. Indexing such spectra has been an important topic in bioinformatics in the past decade. For example, the *k*-spectrum and the associated de Bruijn graph is a central tool in genome assembly [11]. In metagenomics, *k*-spectra have found their place as a useful approximation of the sequence content of the sample, allowing rapid similarity estimation between data collected from sequencing estimates [29, 37]. In applications, typical values for *k* are in the range from 20 to 100.

There are multiple design goals for efficient representations of *k*-mer spectra. In general, the index should be small enough to fit in the main memory of a server machine, while offering fast support for *membership queries*, that is, queries asking whether a given *k*-mer (string of length *k*) is part of the spectrum. Additional query support may include querying for the neighbors of a *k*-mer, that is, *k*-mers that share a suffix or a prefix of length *k*−1 with the current *k*–mer. This allows fast simulation of the de Bruijn graph of the spectrum. The BOSS data structure [6] is a popular solution that meets all the above requirements. Other methods include hashing [32], Bloom filters [38] and the FM-index-based DBGFM structure [9]. Some more recent solutions emphasize the need for dynamic operations, allowing insertion or deletion of data on the index after it has been built [3, 12, 4, 1]. It is also desirable to be able to support attaching some satellite data to each *k*-mer, like is done in, e.g., colored de Bruijn graphs [21, 33, 20, 32, 30, 23], which are now in widespread use. We refer the reader to the recent surveys of Chikhi [8] and Marchet et al. [31] for comprehensive surveys on existing methods.

In this work, we define a static representation of *k*-mer spectra which we call the *Spectral Burrows-Wheeler Transform*, or SBWT for short (we use a capital letter S to disambiguate from the sBWT of Chang et al [7], where the letter s stands for *Schindler*). The SBWT is an evolution of the BOSS data structure [6], which is an indexed representation of the edge-centric de Bruijn graph, based on a version of the Burrows-Wheeler transform. The SBWT differs from the BOSS in that it is node-centric, and more general – the BOSS data structure can be seen as a particular implementation of the SBWT. The SBWT can also be seen as a specialization of the Wheeler graph framework [16] into *k*-spectra, taking full advantage of the properties of the special case.

The SBWT, which we define in Section 3, is a particular sequence of subsets from the alphabet of the input string. To implement *k*-mer membership queries on the SBWT, a form of *rank queries* on subset sequences is required. A subset rank query takes in a character *c* and an index *i*, and returns the count of how many of the first *i* subsets in the sequence of subsets contain *c*. We propose four possible index data structures for subset rank queries, leading to four different SBWT index structures, which we call ConcatSBWT, MatrixSBWT, SplitSBWT and SubsetwtSBWT. ConcatSBWT is a simplified version of the original BOSS representation, whereas the other three are novel variants offering different time-space tradeoffs. SubsetwtSBWT is based on a new data structure we call the *subset wavelet tree*, which is of independent interest. SplitSBWT uses a practical version of subset wavelet tree that is tailored for an SBWT of an input string with a small alphabet, such as the DNA alphabet. MatrixSBWT is a simple variant suitable for small alphabets, that is only slightly larger than the others in practice, but offers extremely fast subset rank queries and *k*-mer search operations.

We then show that it is possible to use entropy coding methods to compress the space of these data structures while retaining query support. In particular, we show that MatrixSBWT implemented with bit vectors compressed to the zeroth order entropy leads to a data structure taking 3.25 bits per *k*-mer on the DNA alphabet, matching the navigational lower bound of Chikhi et al. [9]. An important caveat is that the lower bound of Chikhi et al. is for an arbitrary set of *k*-mers, not for the spectrum of a single string.

The space on a general alphabet of size *σ* is (*n* + *k*)(log *σ* + 1*/* ln 2) + *o*((*n* + *k*)*σ*), where *n* is the number of *k*-mers in the spectrum. The data structure can answer *k*-mer membership queries in *O*(*k*) time, improving on the original BOSS data structure, which takes *O*(*k* log *σ*) time for membership queries.

The index structures SplitSBWT and SubsetwtSBWT are aimed at occupying space that is *lower* than the navigational lower bound. This is achieved by exploiting the uneven distribution of the subsets in the subset sequence of the SBWT. We aim to compress the size of the data structures down to the zeroth order entropy of the *subset sequence*, where each subset is considered as a symbol. We show that this method allows us to get down to 2.44 bits per *k*-mer on an E. coli pangenome.

In practice, our methods lead to a radically new level of performance for succinct de Bruijn graphs, significantly outperforming the best previous approach [33] when space-usage is equated and simultaneously offering a range of attractive space-time tradeoffs. Two highlights are (1) an index that takes only 4 to 5 bits per *k*-mer on our genomic datasets, and is 120 to 242 times faster than the BOSS implementation of VARI and (2) an index taking only 2.6 to 2.8 bits per *k*-mer while being 21 to 56 times faster than VARI.

## 2 Preliminaries

Throughout we will consider a *string S* = *S*[1..*n*] = *S*[1]*S*[2] … *S*[*n*] on an integer alphabet Σ of *σ* symbols. The *colexicographic order* of two strings is the same as the lexicographic order of their reverse strings. The *substring* of *S* that starts at position *i* and ends at position *j, j* ≥ *i*, denoted *S*[*i*..*j*], is the string *S*[*i*]*S*[*i* + 1] … *S*[*j*]. If *i > j*, then *S*[*i*..*j*] is the empty string *ε*. A suffix of *S* is a substring with ending position *j* = *n*, and a prefix is a substring with starting position *i* = 1. We use the term *k*-mer to refer to a (sub)string of length *k*.

A de Bruijn graph (dBG) of order *k* is directed labelled graph built from a set of *k*-mers *S*. There are two prevailing views of a dBG, and we define both here. In the *node-centric* dBG, the node set is given by *S* and there is an edge from node *u* to *v* iff the last *k*−1 symbols of *u* are equal to the first *k* − 1 symbols of *v*. In an *edgecentric* dBG, the node set is given by the set of (*k* − 1)-mers present in *S*, and, for every *x* ∈ *S*, there is an edge from *x*[1..*k* − 1] to *x*[2..*k*]. In other words, the *k*-mers of *S* are nodes in the node-centric dBG and edges in the edge-centric dBG. See Figure 1. Node-centric and edge-centric dBGs represent equivalent information, however, as we shall see, the ease of representing and navigating them can differ significantly.

**Figure 1:**
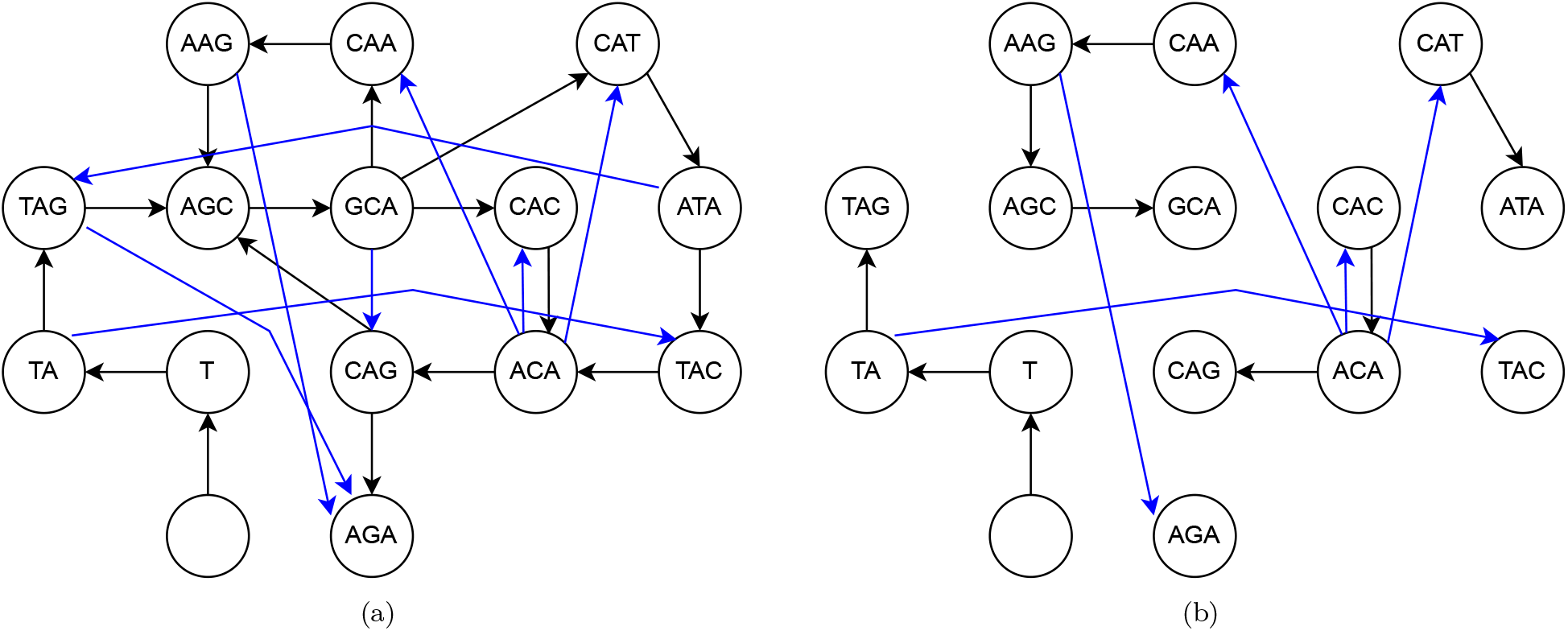
(a) The de Bruijn graph for the string TAGCAAGCACAGCATACAGA. Black edges are in the edge-centric graph. Blue edges are only in the node centric graph; (b) The de Bruijn graph of TAGCAAGCACAGCATACAGA after edge pruning. Every node except for the node of the empty string has exactly one incoming edge, such that the incoming path spells the colexicographically smallest incoming path in the graph prior to pruning.

A key tool in the design of succinct data structures is the support for the *query* operations rank, select, and access, on a bit string *X* of length *n* defined as follows (for *i* ≤ *n* and *x* ∈ {0, 1}):

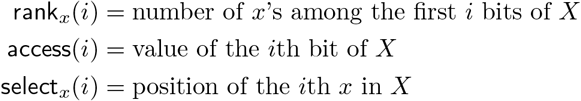

Classical techniques (see, e.g., [34]) require *n* + *o*(*n*) bits to support each of the above queries in *O*(1) time. However, the information theoretic lower bound on space usage for a bit string of length *n* having *n*_1_ 1s,is 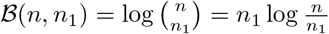 bits.

There are data structures that come within a lower order term of this lower bound while still supporting fast rank, select, and access operations. Perhaps the foremost of these, known as “RRR”, is due to Raman, Raman, and Rao Satti [41] and takes space ***B***(*n, n*_1_) + *o*(*n*) and answers all queries above in *O*(1) time. Fast implementations of RRR have been studied by several authors [35, 18, 24].

Another notable compressed data structure for bit strings is the so-called “Elias-Fano” (or EF) scheme [43, 14, 15], which occupies 2*n*_1_ + *n*_1_⌈log(*n/n*_1_)⌉ bits and supports rank in *O*(log(*n/n*_1_)) time and select_1_(*i*) in *O*(1) time, and tends to be faster than RRR in practice when applied to very sparse bit strings. Like RRR, the efficient implementation of EF has also received considerable practical attention [28, 36].

Rank, access and select queries are also sometimes needed on strings with an alphabet larger than 2. The wavelet tree data structure [19] supports these queries in *O*(*n* log *σ*) bits of space and *O*(log *σ*) time, where *n* is the length of the string and *σ* is the size of the alphabet.

In our analysis later in the paper, we will also make use of the entropy of a probability distribution *p*, which is denoted *H*(*p*) and is defined as:

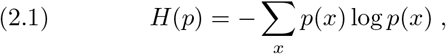

where the sum is over the domain of *p*.

A useful form of the entropy for strings is the so-called zeroth-order empirical entropy, denoted *H*_0_(*S*) for a string *S*, or just *H*_0_ when the context is clear. In particular,

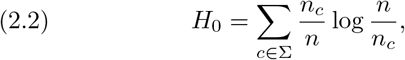

where *n* = |*S*| is the length of *S* and *n*_*c*_ is the number of occurrences of the symbol *c* in *S*. We remark that ***B***(*n, n*_1_) is bounded above by *nH*_0_.

## 3 The Spectral Burrows-Wheeler Transform

In this section we define the Spectral Burrows-Wheeler transform, and the spectrum membership query algorithm based on it. We begin with two basic definitions:

### Definition 1.

*(k-spectrum). The k-spectrum of a string T, denoted with S*_*k*_(*T*), *is the set of all k-mers of the string T*.

### Definition 2.

*(k-prefix set). The k-prefix set of a string T is defined as the left-padded set of prefixes P*_*k*_(*T*) = {$^*k*−*i*^*T* [1..*i*] | *i* = 0, …, *k* − 1}, *where* $ *is a special character not found in the alphabet, that is smaller than all characters of the alphabet*.

For an illustration of the above definition, consider the string *T* = TAGCAAGCACAGCATACAGA, for which we would have *S*_3_(*T*) = AAG, ACA, AGA, AGC, ATA, CAA, CAC, CAG, CAT, GCA, TAC, TAG}, and *P*_3_(*T*) = { $$$, $$T, $TA }.

We are now ready to define the Spectral BWT.

### Definition 3.

*(Spectral BWT, SBWT). Let T be a string from an alphabet* Σ *of size σ. The spectral BWT of order k of T is a mapping from S*_*k*_(*T*) *to a sequence X*_1_, *X*_2_, … *X*_*n*_ *of subsets of* Σ. *The set X*_*i*_ *is defined as follows. Let x*_*i*_ *be the colexicographically i-th k-mer in S*_*k*_(*T*) ∪ *P*_*k*_(*T*). *If x*_*i*_ *is the colexicographically smallest k-mer in S*_*k*_(*T*) ∪ *P*_*k*_(*T*) *that has x*_*i*_[2..*k*] *as a suffix, then X*_*i*_ *is the set of last characters of k-mers y* ∈ *S*_*k*_(*T*) ∪ *P*_*k*_(*T*) *that have x*_*i*_[2..*k*] *as a prefix. Otherwise, x*_*i*_ *is an empty set*.

Continuing the example above, the colexicographically ordered list of *S* _3_ (*T*) ∪ *P* _3_ (*T*) is: $$$, CAA, ACA, GCA, AGA, $TA, ATA, CAC, TAC, AGC, AAG, CAG, TAG, $$T, CAT; and the SBWT is the sequence of sets: {T}, {G}, {ACGT}, ∅, ∅, {CG}, ∅, {A}, ∅, {A}, {AC},∅,∅, { A }, { A }.

The sets in the SBWT represent the labels of outgoing edges in the node-centric de Bruijn graph, such that we only include outgoing edges from *k*-mers that have a different suffix of length *k*−1 than the preceeding *k*-mer in the colexicographically sorted list. The padding of dollar-symbols in Definition 3 is a technical detail that is required to make the SBWT work. Alternatively, the sequence *T* could be made cyclic, and the need for *P*_*k*_(*T*) avoided.

We now describe how to implement efficient *k*-mer membership queries on the spectrum of the input string, using only the information encoded in the SBWT. Here it is beneficial to view the spectrum as a de Bruijn graph. We can think of the nodes as being ordered by the colexicographic order of the corresponding *k*-mers. We define an order for the edges such that the edges are sorted primarily by the edge label, and secondarily by the order of the origins of the edges. We denote with *R*(*e*) the rank of edge *e* in this order. Because the graph is a de Bruijn graph, the indices of the destination nodes of the edges are in the same order as the ranks *R*(*e*), that is, if *R*(*e*_1_) *< R*(*e*_2_), then the destination of edge *e*_1_ is larger than the destination of edge *e*_2_.

Due to Definition 3, every node has exactly one incoming edge, except for the node corresponding to *k*-mer $^*k*^, which has no incoming edge. This is because when there are multiple nodes that could have an edge to the same node, we always use the node corresponding to the colexicographically smallest *k*-mer choice, and since the graph is built from a spectrum of a single string, every node has at least one candidate incoming edge (except for the node of *k*-mer $^*k*^). This means that the destination of edge *e* is the node with index *R*(*e*) +1 in the sorted order.

Thus, the entire graph can be extracted from just the SBWT alone. The *k*-mer label of a node is spelled by any incoming path of length *k* to the node (the properties of de Bruijn graphs ensure that every incoming path of length *k* has the same label). This shows that the SBWT is invertible in the sense that is it possible to extract the original spectrum back from the SBWT.

This graph is also a Wheeler graph [16], which means that the generic Wheeler graph index could be used to index the graph. The Wheeler graph index however requires the storage of the sequence of indegrees and outdegrees of the nodes in the graph, whereas in the SBWT this is not required because every node apart from the node of the empty string has in-degree of exactly 1, and the sequence of outdegrees is already included in the sizes of the sets of the outgoing edge label sets. The most important conceptual advance over the Wheeler graph index, however, is a reformulation of the *k*-mer search problem in terms of *subset rank queries*.

**Subset rank query**: Let *X*_1_, … *X*_*n*_ be a sequence of subsets of an alphabet Σ = {1, …, *σ* }. A subset rank query takes as an input an index *i* and a character *c* ∈ Σ, and returns the number of subsets *X*_*j*_ with *j* ≤ *i* such that *c* ∈ *X*_*j*_.

We now describe a *k*-mer search routine that uses subset rank queries as the only subroutine. The search routine works by searching the *k*-mer character by character from left to right, maintaining the interval of nodes that are suffixed by the prefix that has been processed so far. Suppose we have an interval [*i, j*] of nodes suffixed by prefix *α* of the *k*-mer, and we want to find the interval [*i*′, *j* ′] of nodes suffixed by *αc*, where *c* is the next character in the *k*-mer. This is equivalent to following all edges labeled with *c* from the nodes in [*i, j*]. Due to the way the edges are defined, the end points of these edges are a contiguous range [*i*′, *j*′] such that *i*′ is the destination of the first outgoing edge labeled *c* from [*i*..*j*], and *j*′ is the destination of the last one. Let *C*[*c*] be the number of edge labels with label smaller than *c* in the graph. Now we have the following formulas:

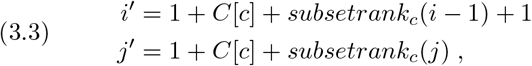

where *subsetrank*_*c*_ is a subset rank query on the sequence of subsets in the SBWT. The “+1” at the start of the formulas is to skip over the node of $^*k*^. The values *C*[*c*] can be precomputed for all characters *c* ∈ Σ. By iterating these formulas *k* times, we have the *k*-mer search routine. See Algorithm 1. This establishes the result below.

### Lemma 3.1.

*The SBWT supports k-mer membership queries in O*(*kt*) *time, where t is the time for a subset rank query*. □

### 3.1 Extension to Multiple Strings

In this subsection, we define an extension of the SBWT to multiple input strings *T*_1_, … *T*_*m*_. The easiest way to extend the SBWT to multiple strings would be to just replace the spectrum *S*_*k*_(*T*) and the *k*-prefix set *P*_*k*_(*T*) in Definition 3 with the unions *S*_*k*_(*T*_1_) ∪ … ∪ *S*_*k*_(*T*_*m*_) and *P*_*k*_(*T*_1_) ∪ … ∪ *P*_*k*_(*T*_*m*_) respectively. However, if the input consists of a large number of short strings, such as DNA sequence reads, this method can introduce a significant space overhead, because every string *T*_*i*_ adds *k* padded prefixes to the union. On the other hand, the prefixes are only there to ensure that every *k*-mer has at least one incoming edge. Therefore, we only need to include the *k*-prefix sets of those strings *T*_*i*_ where the leftmost (*k* − 1)-mer of *T*_*i*_ does not appear as a suffix of any *k*-mer in *T*_1_, …, *T*_*m*_, because otherwise an incoming edge will already be provided by the existing *k*-mer whose suffix is the leftmost (*k* − 1)-mer of this string. Let *R*(*T*_1_, …, *T*_*m*_) be the set of indices *i* such that *T*_*i*_ have this property, and let

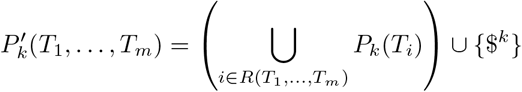

be the modified prefix set. The *k*-mer $^*k*^ is always added for convenience to match the property in the regular SBWT where the *k*-mer $^*k*^ always exists. Finally, let

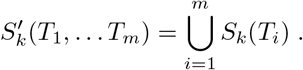

Now, we define the Multi-string Spectral BWT as follows:

#### Definition 4.

*(Multi-SBWT). Let* {*T*_1_, … *T*_*m*_}*be a set of strings from an alphabet* Σ *of size σ and let* 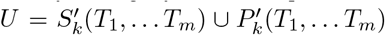. *The multi-SBWT of order k of* {*T*_1_, … *T*_*m*_} *is a sequence X*_1_, *X*_2_, … *X*_*n*_ *of subsets of* Σ. *The set X*_*i*_ *is defined as follows. Let x*_*i*_ *be the colexicographically i-th k-mer in U*. *If x*_*i*_[2..*k*] *is the* Lemma 3.1. *The SBWT supports k-mer membership queries in O*(*kt*) *time, where t is the time for a subset rank query. colexicographically smallest k-mer in U that is suffixed by x*_*i*_[2..*k*], *then X*_*i*_ *is the set of last characters of k-mers y* ∈ *U that are prefixed by x*_*i*_[2..*k*]. *Otherwise, x*_*i*_ *is an empty set*.

All the properties required in the SBWT for the *k*-mer search in Algorithm 1 to work are preserved in this definition, so the *k*-mer search routine still works for the multi-SBWT without modifications.

### 3.2 Streaming search queries

In applications, we often want to search for all *k*-mers in a single long query string. We call this a *streaming query*. If the input query has *m k*-mers, it takes *O*(*mk*) subset rank queries to search them all using Algorithm 1 for each *k*-mer separately. However, it is possible to speed this up to *O*(*m*) subset rank queries in the fortunate case where all of the *k*-mers are found in the index, which often happens in realistic use cases.

#### Algorithm 1 SBWT *k*-mer search query.

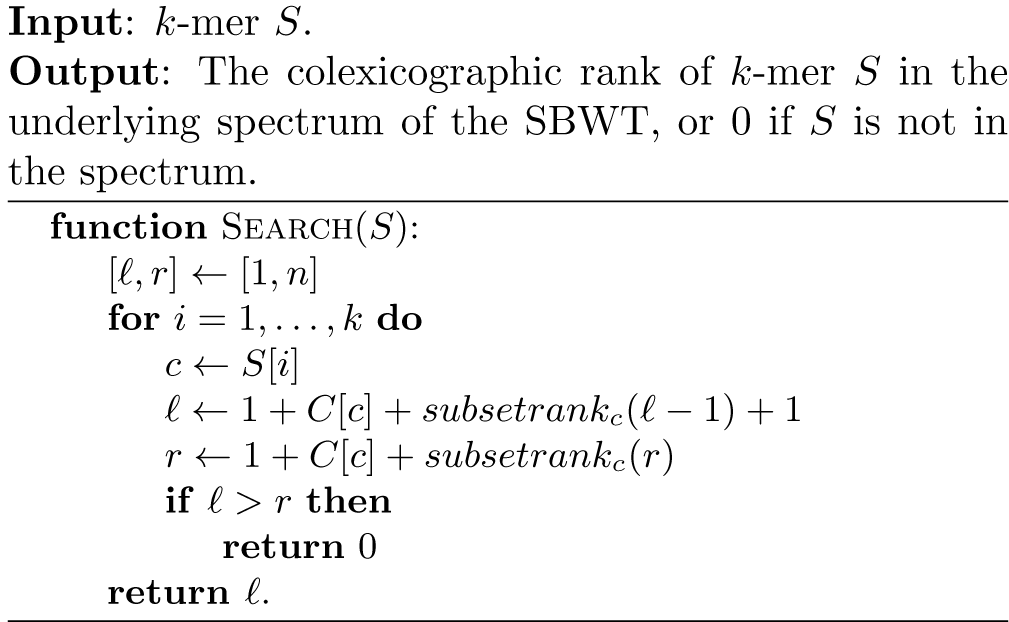

This is made possible by searching the *k*-mers from left to right and reusing computation from the previous iteration. The idea is to build a bit vector that allows us to go from the index of a *k*-mer *x* in the SBWT to the indices of the range of *k*-mers that have *x*[2..*k*] as a suffix, and then execute one more iteration of the loop in Algorithm 1 from this range with the next character in the query. The bit vector marks the first set of the SBWT and all sets where the suffix of length *k*−1 the corresponding *k*-mer is different from the suffix of the previous *k*-mer in the SBWT. In our running example from Definition 3, we would mark the indices of *k*-mers $$$, CAA, ACA, AGA, $TA, CAC, AGC, AAG, $$T and CAT, giving us the bit vector *B* = 111011010110011. Now, if we are at index *i* corresponding to *k*-mer *x*, the range of (*k*−1)-mer *x*[2..*k*] is [pred(*B, i*), succ(*B, i*) − 1], where pred(*B, i*) and succ(*B*,i) are the indices of the previous and next 1-bits respectively in B from index *i* (if *B*[*i*] = 1, then pred(*B, i*) = succ(*B, i*) = *i*). The operations pred and succ can be implemented as a simple for-loop because the gap between consecutive 1-bits in *B* can not be larger than |Σ| + 1. This is because the number of distinct *k*-mers that have the same suffix of length *k*−1 is at most |Σ| + 1, counting in the possibility of a dollar-symbol at the start of the *k*-mer.

This can be seen as a restricted version of the technique used in succinct variable order de Bruijn graphs [5], which allow streaming queries in *O*(*m*) operations, where *m* is the number of consecutive *k*-mers queried, even when none of the *k*-mers are found in the index. This general scheme could also be used in the SBWT, adding an overhead of *O*(log *k*) bits for each set in the SBWT.

### 3.3 Construction

In this section we describe an SBWT construction algorithm that takes as an input a colexicographically sorted set of distinct *k*-mers. Such a sorted database can be efficiently built in parallel by, e.g., using the KMC algorithm [13] to compute the set of distinct *k*-mers of the input, and by sorting them with radix sort.

We describe our construction algorithm in two phases. In the first phase, we find the SBWT sets for all *k*-mers in the input, and report *k*-mers *x*_*i*_ that require the addition of prefixes *P*_*k*_(*x*_*i*_) as in Definition 4. This is done as follows. Denote with *L* = *x*_1_, …, *x*_*n*_ the colexsorted list of input *k*-mers. Let us now focus on *k*-mers that end in a fixed character *c*. We generate the list of *k*-mers *L*_*c*_ = *x*_1_[2..*k*] *c* …, *x*_*n*_[2..*k*] *c*. This list is also in colex-sorted order, and contains all input *k*-mers that end in *c* and have a predecessor (and possibly *k*-mers that are not in the input at all). We thus report those *k*-mers of *L* ending in *c* that do not appear in *L*_*c*_. As both *L* and *L*_*c*_ are sorted, this can be done in linear time by scanning the lists. This process is carried out for all characters *c* of the alphabet. The SBWT sets of the *k*-mers can also be read from the lists *L* and *L*_*c*_: The SBWT set of *k*-mer *x*_*i*_ is empty if *L*[*i*][2..*k*] = *L*[*i* − 1][2..*k*], and otherwise the set contains character *c* iff *L*_*c*_[*i*] was found in *L*. Since the lists are sorted, all this information can be collected in one streaming pass over the lists *L* and *L*_*c*_ for each *c* ∈ Σ. Lists *L* and *L*_*c*_ need not be materialized in memory as a whole. Instead, list *L* can be streamed from disk and the lists *L*_*c*_ generated on the fly. See the pseudocode in Algorithm 3 in Appendix D for more details.

Next, we add the prefix sets *P*_*k*_(*x*_*i*_) for the *k*-mers *x*_*i*_ that lacked a predecessor in the first phase. We start by generating the prefix set *P*_*k*_(*x*_*i*_) for all such *k*-mers *x*_*i*_. Let us denote the resulting list with *L*′. We attach to each generated prefix the character that follows that prefix in the original *k*-mer. We then sort the prefixes, merging duplicates such that we collect the associated characters in each group of duplicates. The collected sets are the SBWT sets of the corresponding prefixes. In typical applications, the total size of the prefix sets tends to be small, so the sorting can usually be done in internal memory – if not, we may have to fall back to a disk-based sorting algorithm. Finally, we merge the sorted list of SBWT sets of *L* with the sorted list SBWT of sets of *L*′ to form a single sorted list containing the SBWT sets of all *k*-mers and required prefixes. See Figure 6 for an example of the whole process including phases 1 and 2.

## 4 Data Structures for Subset Rank Queries

In this section, we propose four different succinct data structures for subset rank queries. Our treatment here is not intended to be exhaustive, but rather to demonstrate that an interesting range of space-time tradeoffs are possible. We use the notation from the previous section, that is, we are indexing a sequence *X*_1_, … *X*_*n*_ of subsets of an alphabet Σ = {1, …, *σ*}. A subset rank query takes as an input an index *i* and a character *c* ∈ Σ, and returns the number of subsets *X*_*j*_ with *j* ≤ *i* such that *c* ∈ *X*_*j*_.

### 4.1 Plain Matrix Representation

This data structure uses a binary matrix *M* of size *σ* ×*n*, such that *M* [*i*][*j*] = 1 iff subset *X*_*j*_ contains the *i*-th character in the alphabet. See Figure 2. The rows of the matrix are indexed for constant-time succinct rank queries. The subset rank query for the *i*-th character of the alphabet up to index *j* is answered in contant time with a rank query on row *M* [*i*] up to index *j*.

**Figure 2:**
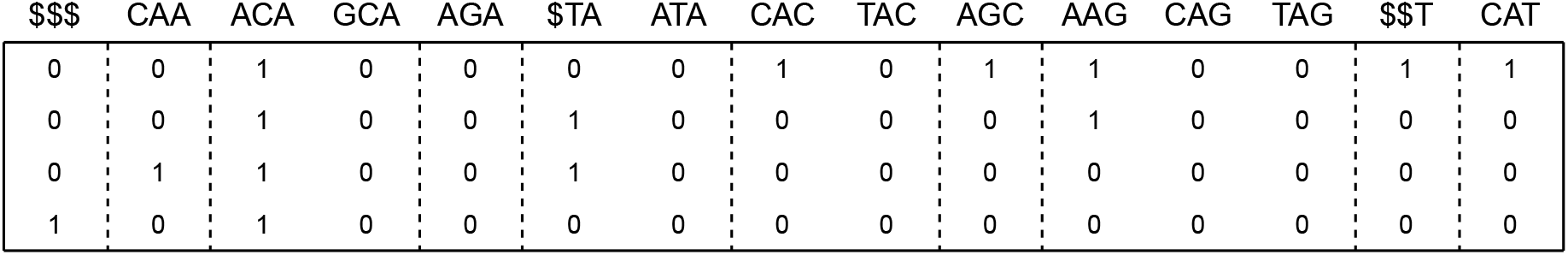
MatrixSBWT of TAGCAAGCACAGCATACAGA with *k* = 3. The dashed lines indicate borders of suffix groups. Two adjacent columns are in the same group if they have the same suffix of length *k* −1. Bits may be moved horizontally inside a suffix group without affecting the *k*-mer set encoded in the matrix.

### 4.2 Split Representation

This is a version of the plain matrix representation, tailored for the use case of subset rank queries in the SBWT, exploiting the property that, in many use cases in genomics, most of the sets in the SBWT are singletons. Let *M* ^−^ be the submatrix of matrix *M* in the plain matrix representation that contains only the columns of *M* with exactly one 1-bit set, and let *M* ^+^ be the submatrix of *M* containing the rest of the columns. Let *B* be a bit vector of length equal to the number of columns in *M*, marking with 1-bits which columns of *M* are in *M* ^+^.

We index both *M* ^+^ and *M* ^−^ for character rank queries. Matrix *M* ^+^ is indexed like in the plain matrix representation. Matrix *M* ^−^ on the other hand is replaced by a string *W* that is the concatenation of the labels corresponding to the bits in the columns. The string *W* is indexed as a wavelet tree. The bit vector *B* is indexed for rank queries. In summary, the final index consists of just *B, M* ^+^ and *W* and their rank support structures. See Figure 3.

**Figure 3:**
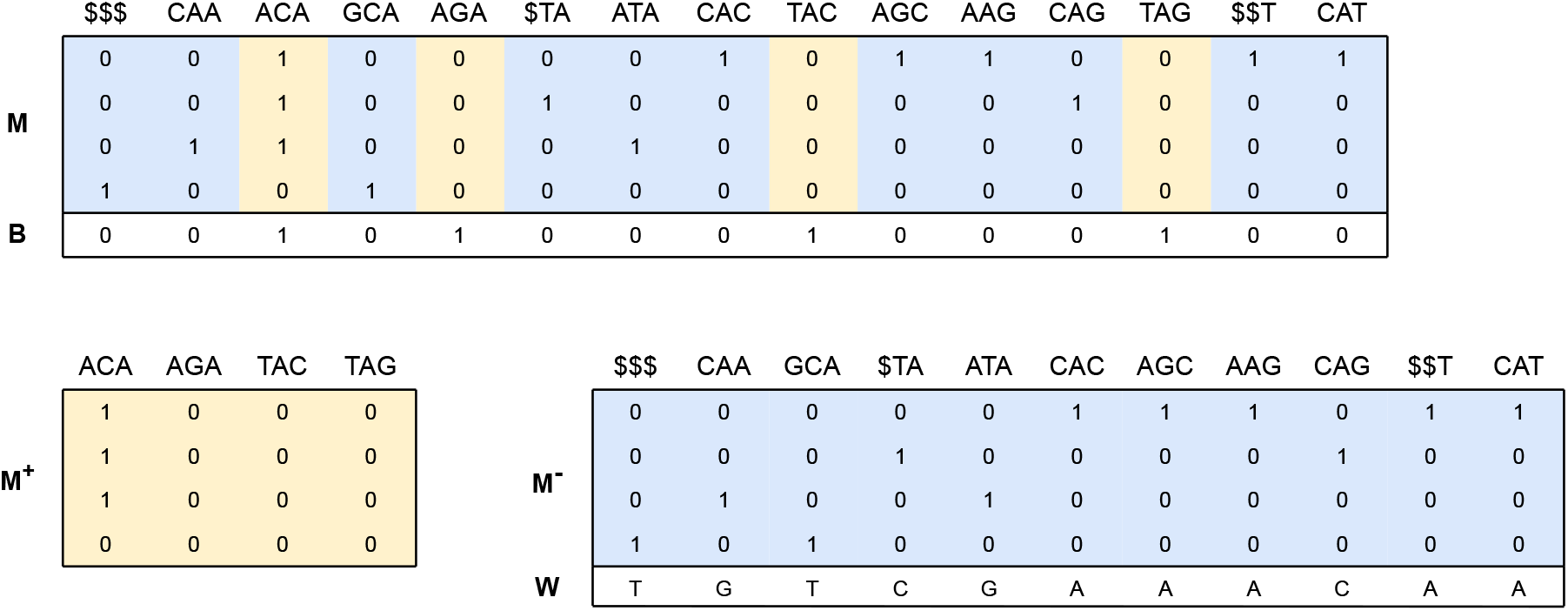
SplitSBWT of TAGCAAGCACAGCATACAGA. The bits are spread out inside a suffix group to maximize the number of singleton sets. The index consists of just *B, M* ^+^ and *W*.

Subset rank query for *i*-th character up to index *j* in the original subset sequence can now be answered by using a rank query on *B* to determine how many of the first *j* columns of *M* went to *M* ^+^ and *M* ^−^, then using the rank structures of *M* ^+^ and *M* ^−^ to count the number of characters in the corresponding prefixes of columns in both matrices, and returning the sum of the two counts. That is, if *r* = *rank*_1_(*B, j*), then answer to the query is *rank*_1_(*M* ^+^[*i*], *r*) + *rank*_*i*_(*W, j* − *r*).

### 4.3 Concatenated Representation

This data structure uses a concatenation of the contents of the subsets, and an encoding of the sequence of sizes of the subsets. In more detail, let *S*(*X*_*i*_) be the concatenation of the characters in subset *X*_*i*_. If *X*_*i*_ is the empty set, we define *S*(*X*_*i*_) = $. We build the string *L* = *S*(*X*_1_)*S*(*X*_2_) … *S*(*X*_*n*_) and index it for rank queries. The empty sets are represented as dollars to be able to encode the sizes of the subsets with a bit vector *B* that is the concatenation 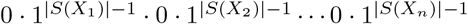. See Figure 4. If the sequence of subsets contains a large number of singleton or empty sets, the bit vector *B* is sparse, and is efficiently compressed using the Elias-Fano encoding. The bit vector *B* is indexed for queries for select-0. A subset rank query for a character *c* up to index *i* is then answered by computing *rank*_*c*_(*L, select*_0_(*B, i* + 1) − 1).

**Figure 4:**
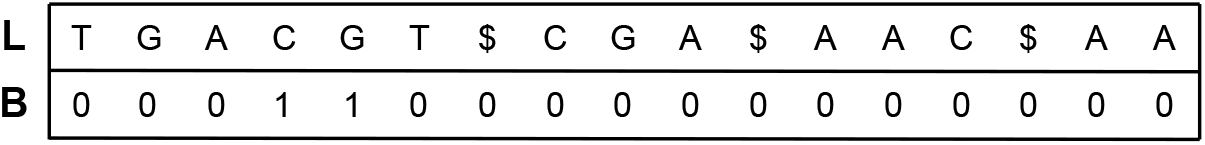
ConcatSBWT of TAGCAAGCACAGCATACAGA. The bits are spread out inside a suffix group to maximize the number of singleton sets.

### 4.4 Subset Wavelet Tree

We build a tree with log *σ* levels (assume for simplicity that *σ* is a power of 2). Each node of the tree corresponds to a part of the alphabet, defined as follows. We denote with *A*_*v*_ the alphabet of node *v*. The root node corresponds to the full alphabet. The alphabets of the rest of the nodes are defined recursively such that the left child of a node *v* corresponds to the first half of *A*_*v*_, and the right child corresponds to the second half of *A*_*v*_. Let *Q*_*v*_ be the subsequence of subsets that contain at least one character from *A*_*v*_. As a special case, the subsequence *Q*_*v*_ also includes the empty sets if *v* is the root.

Each node *v* contains two bit vectors *L*_*v*_ and *R*_*v*_ of length |*Q*_*v*_|. We have *L*_*v*_[*i*] = 1 iff subset *Q*_*v*_[*i*] contains a character from the first half of *A*_*v*_, and correspondingly *R*_*v*_[*i*] = 1 if *Q*_*v*_[*i*] contains a character from the second half of *A*_*v*_. See Figure 5. The bit vectors *L*_*v*_ and *R*_*v*_ may be entropy-compressed efficiently by considering them together as a string from alphabet {0, 1, 2, 3}, such that the *i*-th character is defined as (2 *L*_*v*_[*i*] + *R*_*v*_[*i*]). This will take advantage of the fact that in the case of the SBWT, it is rare that *L*_*v*_[*i*] = *R*_*v*_[*i*] because most of the sets in the SBWT tend to be singletons. Rank queries on *L*_*v*_ can then be implemented by summing the ranks of characters 0 and 2, and rank queries on *R*_*v*_ can be implemented by summing the ranks of characters 1 and 3.

**Figure 5:**
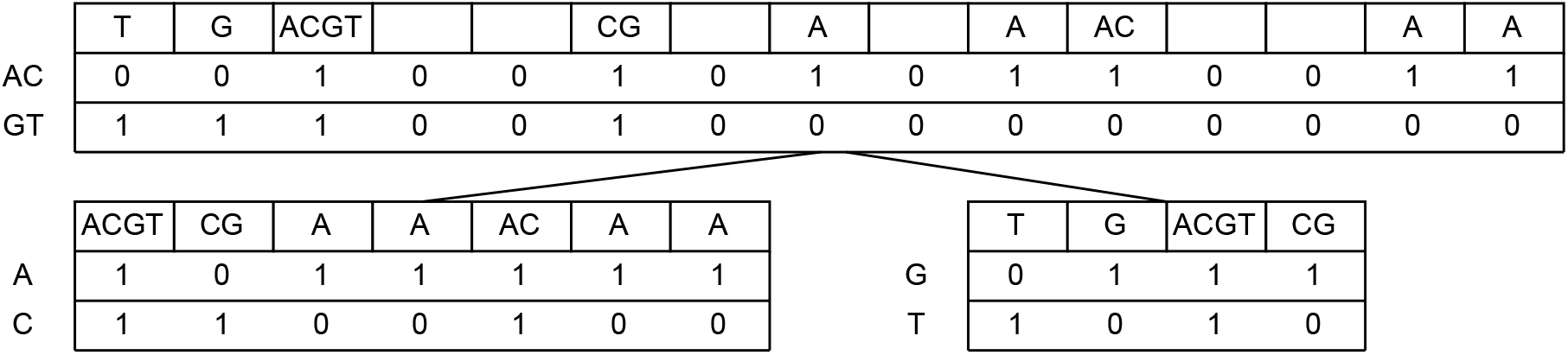
SubsetwtSBWT of TAGCAAGCACAGCATACAGA. The labels are concentrated to single sets inside a suffix group to create many empty sets. The index consists of just the bit vectors in the figure.

To answer our query for a character *c* and position *i*, we traverse from the root to the leaf of the tree that where *A*_*v*_ is the singleton subset *c*. While traversing, we compute for each visited node *v* the length of the prefix in the current subset sequence *Q*_*v*_ that contains all the subsets of *X*_1_, … *X*_*i*_ that have at least one character from *A*_*v*_. This is done by using rank queries on the bit vectors *L*_*v*_ and *R*_*v*_, analogous to a regular wavelet tree query. Pseudocode is given in Algorithm 2.

Query time for the subset wavelet tree is clearly *O*(log *σ*), as constant time is spent at each of the log *σ* levels. For a general sequence of sets, the data structure requires 2*nσ* + *o*(*nσ*) bits of space, as if all sets are full, then each set goes both ways at each level. For the SBWT matrix however, less space is needed. In particular, because each column of the matrix contains exactly one 1-bit, sets can participate in at most one node at each level of the subset wavelet tree, making the number of bits in the bitvectors over all log *σ* levels of the tree 2*n* log *σ*. We thus have the following theorem.

#### Theorem 4.1.

*The subset wavelet tree of the SBWT takes* 2*n* log *σ* + *o*(*n* log *σ*) *bits of space and supports subset rank queries in O*(log *σ*) *time*.

#### Algorithm 2 Subset wavelet tree query.

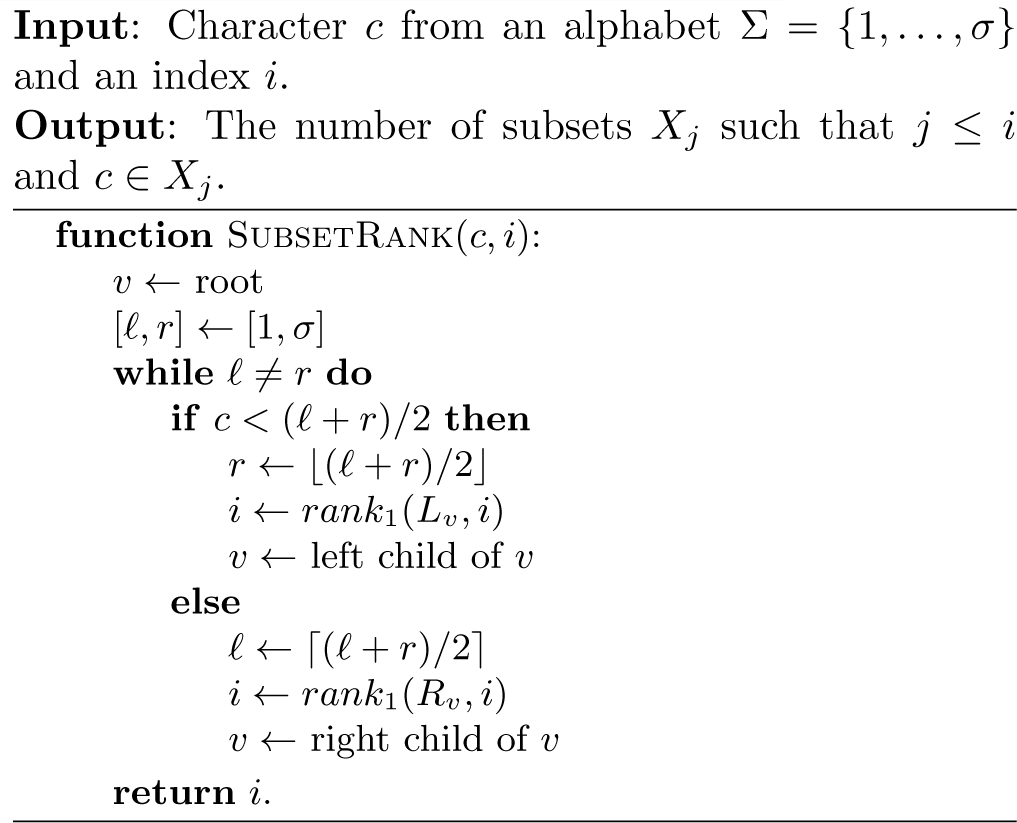

## 5 Analysis

In this section, we prove bounds on the sizes of entropy-compressed versions of the plain matrix representation of an SBWT. The size bounds are expressed in terms of the number of columns of the matrix, which is equal to |*S*_*k*_(*T*)| + *k*, where *T* is the original string.

### Theorem 5.1.

*A plain matrix SBWT representation can be encoded in n*(log *σ* +1*/* ln 2) + *o*(*nσ*) *bits of space, with support for k-mer membership queries in O*(*k*) *time, where n is the number of columns in the matrix, and σ >* 1 *is the size of the alphabet*.

*Proof*. The bit matrix has *σ* rows and *n* columns, and always has *n* − 1 ones. In other words, the fraction of one-bits in the matrix is (*n* −1)*/*(*nσ*), which is less than 1*/σ*. Plugging *Pr*(1) = 1*/σ* and *Pr*(0) = 1 −1*/σ* to the entropy formula (Eq. (2.1)), which is a monotonically increasing function in the interval [0, 1*/σ*], gives

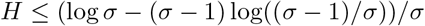

bits of entropy per matrix element, or

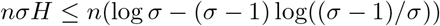

bits of entropy for the whole matrix. The term (*σ* − 1) log((*σ* − 1)*/σ*) is upper bounded by 1*/* ln 2, when *σ >* 1, where ln is the natural logarithm. This can be shown by plugging in *x* = 1*/*(*σ* − 1) to the well-known inequality ln(*x* + 1) ≤ *x*, which gives ln(1*/*(*σ* − 1) + 1) ≤ 1*/*(*σ* − 1), which is equivalent to − (*σ* − 1) log((*σ* − 1)*/σ*) ≤ 1*/* ln 2. So we have:

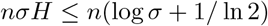

This shows that entropy compressing the matrix to the zeroth order entropy of the bits results in space *n*(log *σ* + 1*/* ln 2). In practice this can be done using the RRR bit vector encoding, which also supports constant-time rank queries on the bit vector with an overhead of *o*(*nσ*). Combining Lemma 3.1 with the RRR encoding gives the result claimed in the theorem.

In the case of the DNA alphabet (*σ* = 4), the space per *k*-mer is (−0.25 log(0.25) − 0.75 log(0.75)) · 4≈3.245 bits ignoring the lower order term. We note that this exactly matches Chikhi’s navigational lower bound of 8− 3 log 3 = 3.245 bits per *k*-mer [9]. This is remarkable, as our data structure is a *membership structure*, whereas the lower bound of Chikhi et al. is for a weaker navigational structure. On the other hand Chikhi’s bound allows an arbitrary *k*-mer set, whereas ours is restricted to the spectrum of a single string.

A different bit vector representation leads to the following space-time tradeoff:

### Theorem 5.2.

*The plain matrix SBWT representation can be encoded in n* ⌈log *σ*⌉ + 2*n bits of space, with support for k-mer membership queries in O*(*k* log *σ*) *time, where n is the number of columns in the matrix, and σ >* 1 *is the size of the alphabet*.

*Proof*. We concatenate the rows of the matrix and represent the resulting bit string, which is of length *nσ* and contains at most *n* 1-bits, using the Elias-Fano scheme. The space required is *n* ⌈log(*nσ/n*) ⌉ + 2*n* = *n* ⌈log *σ*⌉ + 2*n* bits. The time for a single rank query is *O*(log(*nσ/n*)) = *O*(log *σ*), leading to the *O*(*k* log *σ*) time for a *k*-mer membership query as claimed. □

## 6. Experiments

In our experiments, we measure query time and the size of the SBWT using the four different subset rank implementations described in Section 4. We used the value *k* = 31 in all experiments. For each implementation, we include a variant that has entropy compression, and a variant that does not. The entropy compression is done either with RRR bit vectors [41], or the Elias-Fano encoding [43, 14, 15]. We make use of the Succinct Data Structures Library (SDSL) [17] for plain and RRR bitvectors, and use the Elias-Fano bitvector implementation from [28].

We compare our implementations to the BOSS implementation in VARI^1^ [33], and two state-of-the-art hashing-based solutions Bifrost^2^ [20] and Sshash^3^ [39]. We modified all competing tools by adding a timer that starts when a query starts and ends when the query returns. This is to discount for external factors such as disk I/O and the time to parse the queries from FASTA or FASTQ format.

We considered including the early BOSS-like index structure of Rødland [42], which is written in Java. Rødland’s data structure is somewhat similar to our plain matrix variant, but with a different memory layout and an additional bit vector. However, the implementation was unable to process our datasets, probably due to its internal use of 32-bit integers limiting the maximum input size.

Experiments were run on Ubuntu 18.04.5 LTS kernel version 4.15.0-147-generic. The compiler was g++ 9.3.0. The CPU was an Intel Core i7-9700K CPU clocked at 3.60 GHz with with L1d, LIi, L2 and L3 caches of size 256KiB, 256KiB, 2MiB and 12MiB, respectively. The system had 64GiB of DDR4 2400 MHz memory and a 1TB Samsung 860 EVO in the SATA3 bus. The C++ code of the SBWT variants are available at https://github.com/algbio/SBWT. The experiment scripts and the modified versions of all competing tools with our timing code added are available at https://github.com/jnalanko/SBWT_experiments.

### 6.1 Datasets

We experiment on three different data sets that represent typical targets for *k*-mer indexing in bioinformatics applications:

1. A pangenome of 3682 E. coli genomes aiming to replicate the E. coli dataset in the experiments of VARI [33]. The data was downloaded during the year 2020 by selecting a subset of 3682 assemblies listed in ftp://ftp.ncbi.nlm.nih.gov/genomes/ genbank/bacteria/assembly_summary.txt with the organism name “Escherichia coli” with date before March 22, 2016. The resulting collection of genomes is available for download at zenodo.org/ record/6577997.
2. A set of 17,336,887 Illumina HiSeq 2500 sequence reads of length 502 sampled from the human gut (SRA identifier ERR5035349) in a study on irritable bowel syndrome and bile acid malabsorption [22].
3. A set of 1,234,695 genomes of the SARS-CoV-2 virus downloaded from NCBI datasets.

We then construct the multi-string variant of the SBWT defined in Section 3.1 for each of the datasets. K-mers that contain characters outside of the DNA alphabet are discarded. Table 1 shows a number of key statistics for the datasets.

**Table 1:** Key statistics on the three experimental datasets used. The number of unique 31-molecules means the number of 31-mers such that if a *k*-mer occurs both in forward and reverse complement orientations, only one orientation is counted. The number of sequences in the case of E. coli is not the number of genomes, which is 3682, but the number of contigs in assemblies of those genomes. The column UC (unitig count) is the number of unitigs and AUL is the average unitig length. The reported unitigs are the bidirected unitigs computed by BCALM2.

|  | Sequences | Base pairs | Unique 31-mers | Unique 31-molecules | UC | AUL |
| --- | --- | --- | --- | --- | --- | --- |
| **E. coli** | 745,409 | 18,957,578,183 | 253,718,096 | 170,648,610 | 8,939,732 | 49.1 |
| **SARS-CoV-2** | 1,234,695 | 36,808,137,972 | 2,407,721 | 2,407,455 | 178,758 | 43.4 |
| **Metagenome** | 17,336,887 | 8,703,117,274 | 2,761,523,935 | 2,488,344,054 | 105,908,119 | 52.4 |

### 6.2 Index construction

We construct the SBWT by using KMC3 [27] to extract the distinct *k*-mers of the data and sort them. Since KMC3 sorts in lexicographic order, we reverse the sequences before feeding them to the KMC algorithm, and reverse the *k*-mers again when reading the sorted *k*-mer database, to get the *k*-mers in colexicographic order. The memory usage of KMC3 is controllable via a command-line parameter. We used 5GB for the SARS-CoV-2 and E. coli data, and 9GB for the metagenome data. We chose these values to approximately match the memory usage of Sshash. Using more memory would speed up our construction. VARI was constructed using from KMC database using the instructions in the repository, with the exception that we needed to add the flag -cs2 to KMC to prevent VARI from discarding very frequent *k*-mers. The time taken by KMC is included in the total time.

The tools Sshash and Bifrost define a *k*-mer so that a *k*-mer is considered to be equal to its reverse complement. That is, these tools index not the set of *k*-mers, but the set of *k-molecules* of the input, where a single *k*-molecule represents both directions of a *k*-mer. We could mimic this behavious by adding the reverse complements of the input sequences to the input data for SBWT. This way, a *k*-mer membership query in the SBWT returns a positive answer iff the *k*-molecule is found in the input data in either orientation, matching the behaviour of the other two tools. This costs us extra space however, so we build the SBWT indices *without* the added reverse complements. In this case, the query becomes sensitive to the orientation of the *k*-mer, which could be a useful feature in some applications where the orientation is important, such as when designing DNA probes where we want to avoid inadvertent probe-to-probe binding [2]. In any case, even if the index is sensitive to orientation, it is always possible implement queries for *k*-molecules by running the query in both orientations, doubling the query time. VARI always includes both orientations in the index and this aspect is outside of the control of the user.

The Sshash tool requires as input a set of strings where each *k*-molecule occurs only once. The suggested way [39] to construct such a representation is to extract the *unitigs* (non-branching paths in the bidirected de Bruijn graph) of the input using a tool such as BCALM2 [10] or Cuttlefish [26, 25], and stitch unitigs together using UST [40]. We chose BCALM2 over Cuttlefish following the instructions in the Github repository of Sshash. We include the time and memory taken by these tools in the total resource requirements of Sshash. We could also have used stitched unitigs as input to SBWT and Bifrost, but we chose not to do that because one of the advantages of the SBWT construction algorithm is that it works directly on the *k*-mers without a need to extract the unitigs, and because Bifrost has already has a built-in in-memory unitig extraction algorithm that is used in its construction.

Sshash requires the user to choose the value for the length of *k*-mer minimizers used in the data structure. In our experiments, we use the value of 16, which is a reasonable compromise between time and space. In Bifrost, the length of the minimizers are chosen automatically based on the data. For Bifrost we disable the *k*-mer coloring feature to avoid unnecessary computation. The number of parallel threads for all tools was 8, which is equal to the number of cores in the CPU of the test machine.

The SARS-CoV-2 data is peculiar in the sense that the number of distinct *k*-mers in the data is tiny compared to the total length of the input sequences. This means that the extraction of the distinct *k*-mers for SBWT and VARI and unitigs for Sshash and Bifrost dominate the time. For example, extracting the unitigs with BCALM2 took 10min 26s, producing only 24MB of unitigs, from which the Sshash index could be built in just 0.3 seconds. Running KMC on the data for VARI took 6min 30s, resulting in only 2,407,455 *k*-molecules, after which VARI could be built in only 3.45 seconds. The index structures for all tools took space ranging from a few megabytes to tens of megabytes of space.

The datasets of the E. coli pangenome and the metagenome resulted in larger indexes, which gives a better idea of the performance of the construction pipelines after the *k*-mer preprocessing stages. For example, extracting the unitigs with BCALM2 took 56 seconds on E. coli and 12 minutes on the metagenome, which now represents just 10.5% and 19.6% of the total construction time of Sshash. Extracting the k-mers with KMC for VARI took 2min 25s and 1min 16s, which represents 46% and 3% of the total construction time of VARI. We also observed that VARI required on the order of 300GB of temporary disk space during construction, so the disk usage may become prohibitive in larger datasets. Overall, all construction pipelines performed well and finished within approximately an hour on both datasets. For SBWT, extracting the distinct k-mers accounted over 99% the total time in the SARS-Cov-2 datasets, but only for approximately 50% of the total time on E. coli, and only 5% on the metagenome.

The smallest index of all was either the RRR-compressed subset wavelet tree or the Elias-Fano-compressed concatenated representation. The largest index by far was Bifrost, but this is due to Bifrost serializing the index as the set of unitigs of the data in an inefficient ASCII format. On the other hand, the index format of Bifrost does not include the minimizer hash table of the index, which is regenerated every time the index is used for queries, which makes the memory usage during queries even higher than the size on disk.

Full data tables of the results are in the Appendix.

### 6.3 Queries

The query time of all methods depends on whether the *k*-mer is found in the index (a *positive* query) or not (a *negative* query). For the query benchmark, we generated one million positive queries by randomly sampling *k*-mers from the input data, and one million negative queries by sampling *k*-mers uniformly at random from the space of all possible *k*-mers (since *k* = 31, the probability that a random *k*-mer occurs in the index is very small). We also generated longer sequences to test the performance of the tools on streaming queries, where we query all the *k*-mers present in a longer sequence in the order they appear in that sequence^4^. For these queries, as above, we generated both positive and negative sequences by sampling a total of 1 million characters split into sequences of length 500 either from the input sequences (positive), or by drawing the sequences uniformly at random (negative).

Streaming queries employed the approach of Section 3.2, which introduces an overhead of 1 bit per SBWT set, but speeds up queries greatly in the positive case. Numbers used to generate plots for the query experiments are given as tables in Appendices B and C

The indexes for SARS-CoV-2 mostly fit within the L3 cache of the experiment machine, making the results not representative of practical performance on datasets with a larger number of *k*-mers. The peak RAM usage during query time was also dominated by the space to store the queries in a memory buffer. Due to this, we advise the reader to not pay much attention to these results. The tables and plots are included for completeness nonetheless.

On E. coli and the metagenomic read set, the results show that query time of the matrix variant of SBWT is competitive with the state-of-the-art hashing-based approaches. The best case scenario for the SBWT variants is the case of streaming positive queries, where the SBWT requires only two subset rank queries per character in the reads. In this case, the query times are only 110-140 nanoseconds per *k*-mer, beating both SSHash and Bifrost. In the single queries, Bifrost was the fastest, but also took an order of magnitude space more than the competitors. For SBWT, the negative single queries were observed to be roughly twice as fast as the positive queries, as the search can exit early when the search interval becomes empty. This difference will be more pronounced the larger the value of *k* is. In Bifrost, a similar two-fold speedup was observed in negative queries.

Figure 8 shows time-space plots of the query performance of the benchmarked methods on the metagenome data set (the plots for E. coli are similar, see Appendix C). When the streaming support bit vectors is not included (single queries), the plots show that the RRR-matrix variant takes space close to the navigational lower bound of Chikhi et al, as predicted by Theorem 5.1. However, the RRR-matrix variant was dominated in the time-space plane by other SBWT variants. In most cases, the Pareto frontier includes the variants of plain matrix, EF matrix, all split variants, the concatenated EF variant and the RRR subset wavelet tree. Depending on the query type, Bifrost and SShash often also lie on the Pareto frontier. The query time of VARI was the slowest on every query type, which is likely due lack of optimization in the implementation of VARI. The general approach of VARI is better embodied in the concatenated EF variant, which is shown to be comparable to the RRR subset wavelet tree in terms of time and space.

**Figure 6:**
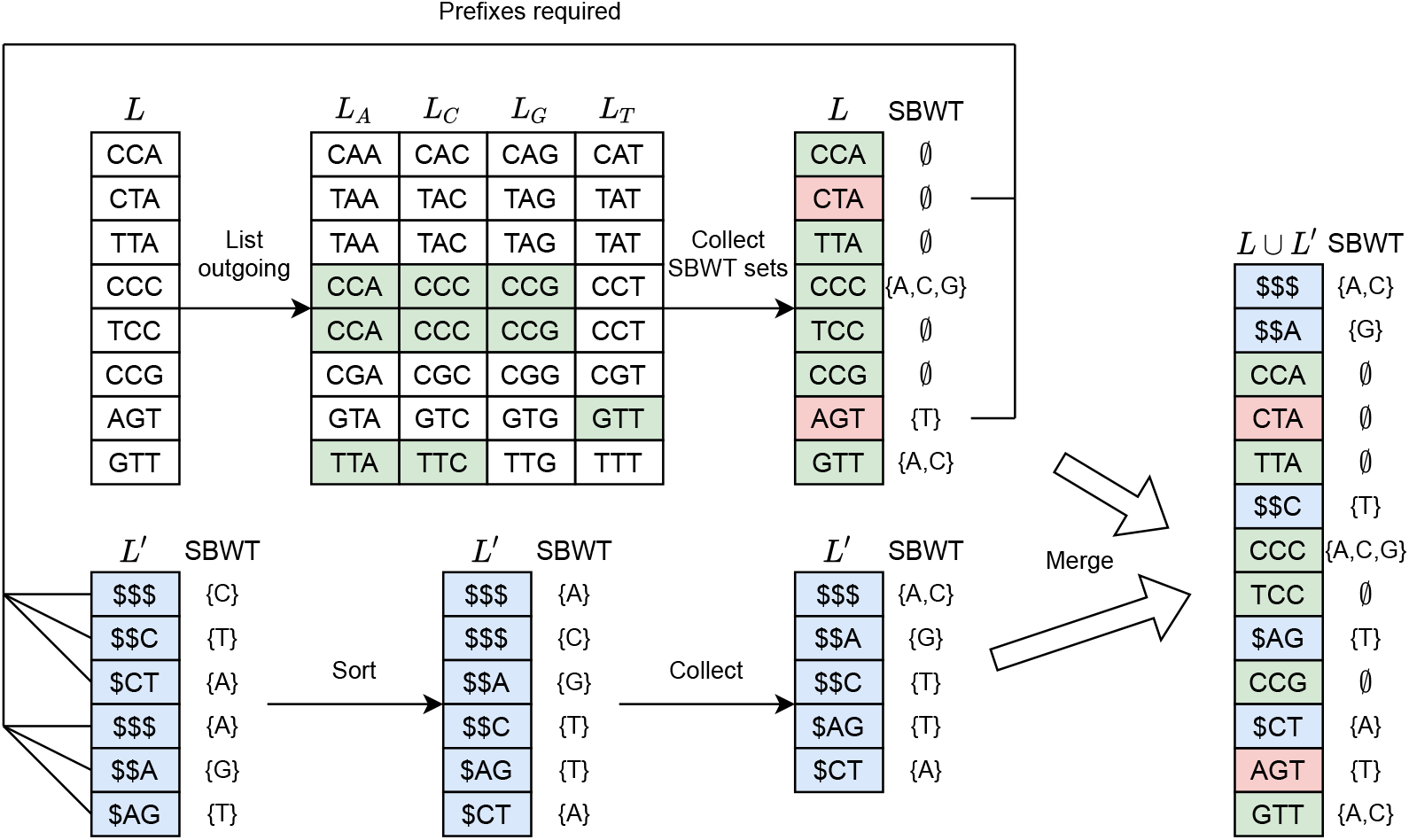
Multi-SBWT construction from a colex-sorted set of *k*-mers CCA, CTA, TTA, CCC, TCC, CCG, AGT and GTT. The *k*-mers marked in green are those who were found to have a predecessor in the graph. The *k*-mers marked in red require the addition of their prefix sets, which are sorted in blue below.

**Figure 7:**
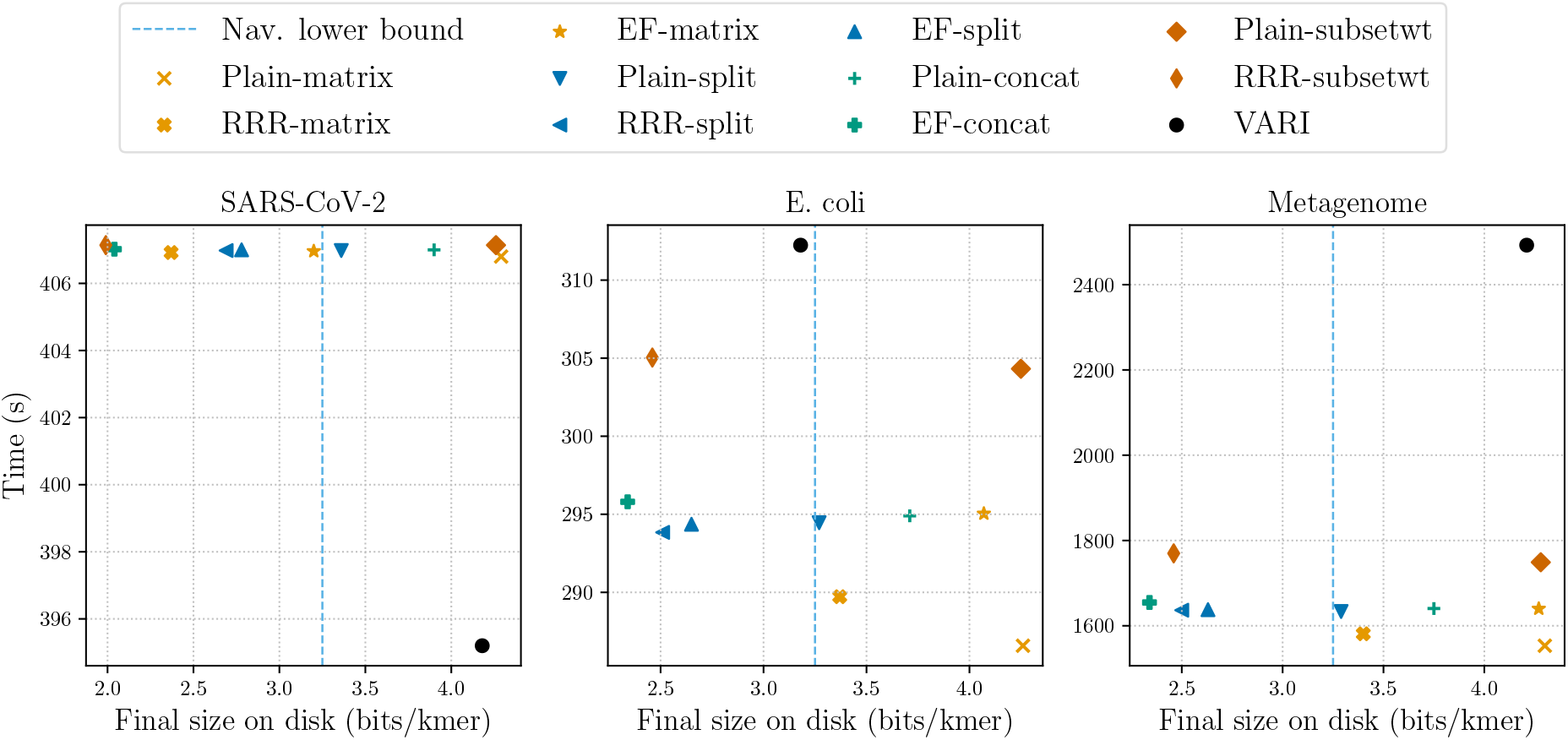
Construction time and final disk space. Tables of the data behind the plots are in Appendix A.

**Figure 8:**
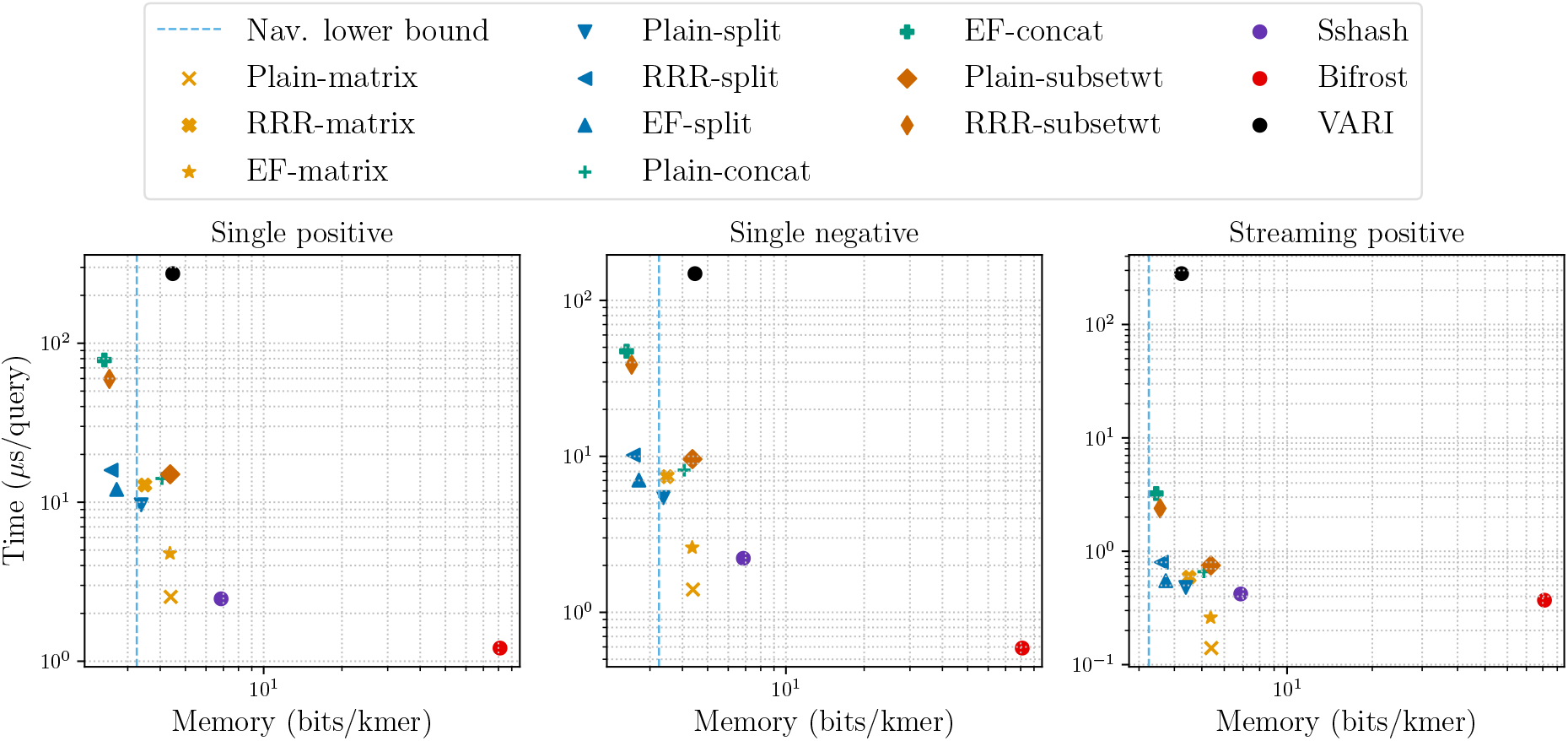
Query results on the metagenome dataset. Streaming negative queries are not plotted due to save space and because streaming negative queries do not give any benefit over single negative queries in the SBWT. See Appendixes B and C for the full data and plots.

**Figure 9:**
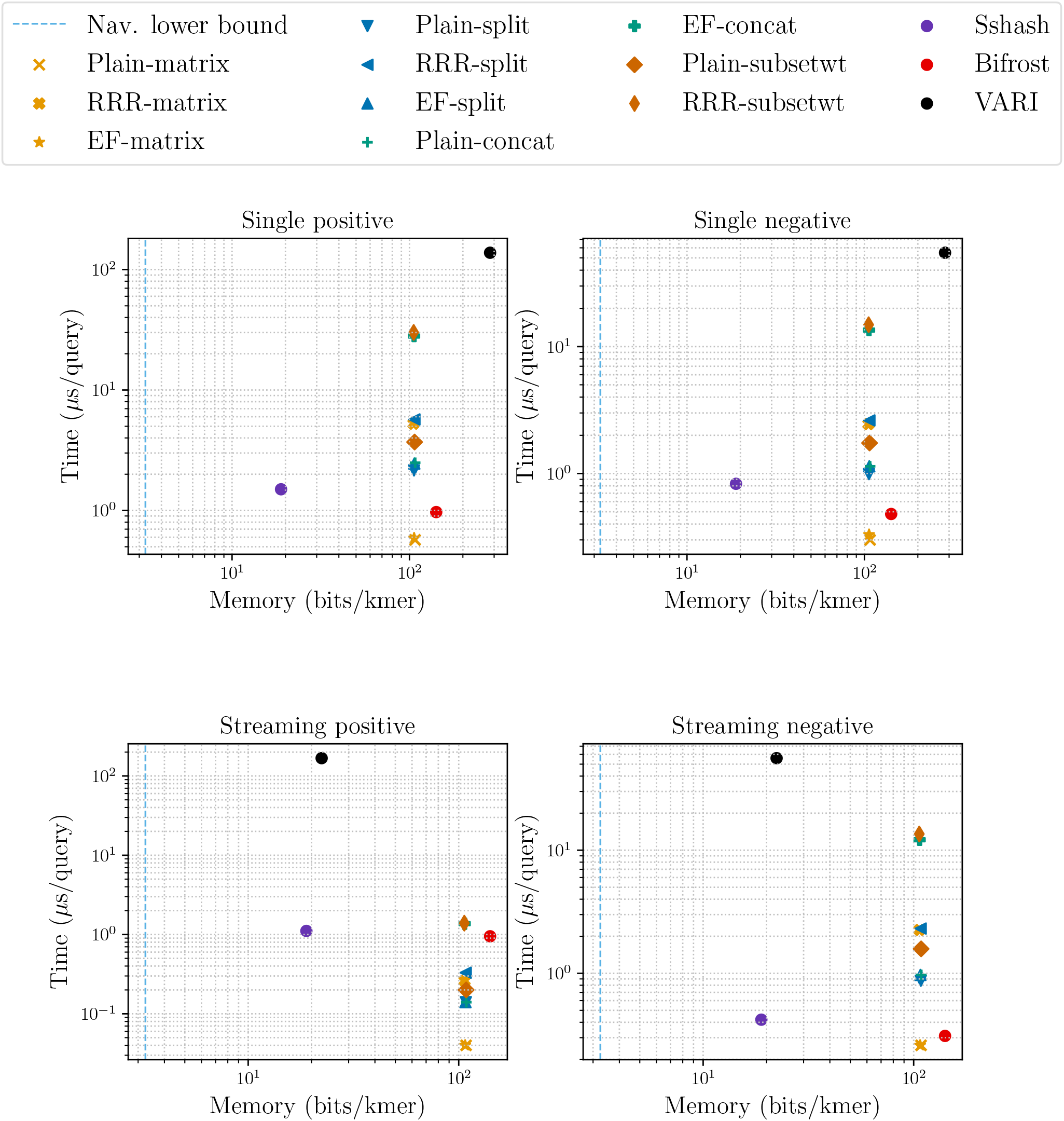
Queries on SARS-CoV-2.

**Figure 10:**
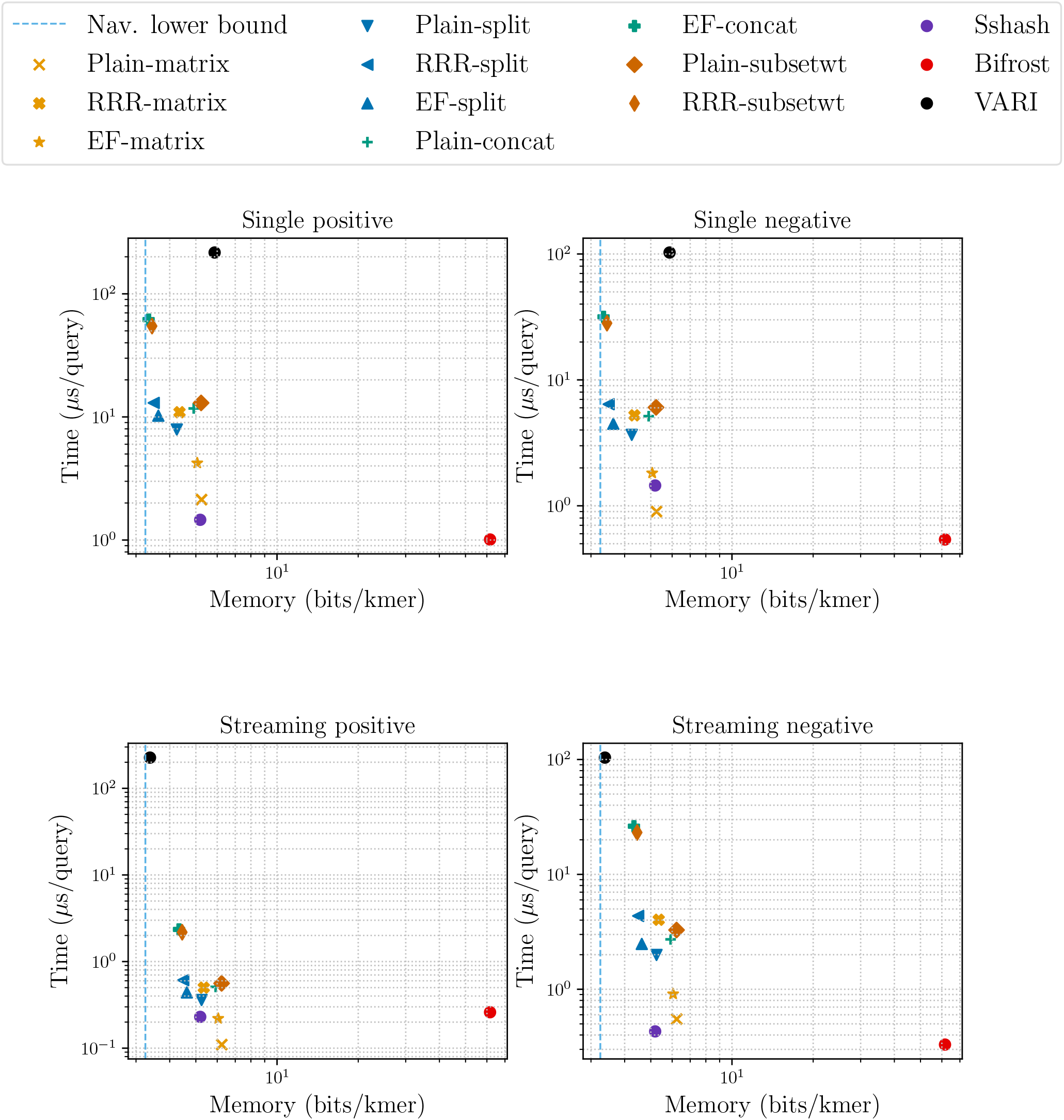
Queries on E. coli.

**Figure 11:**
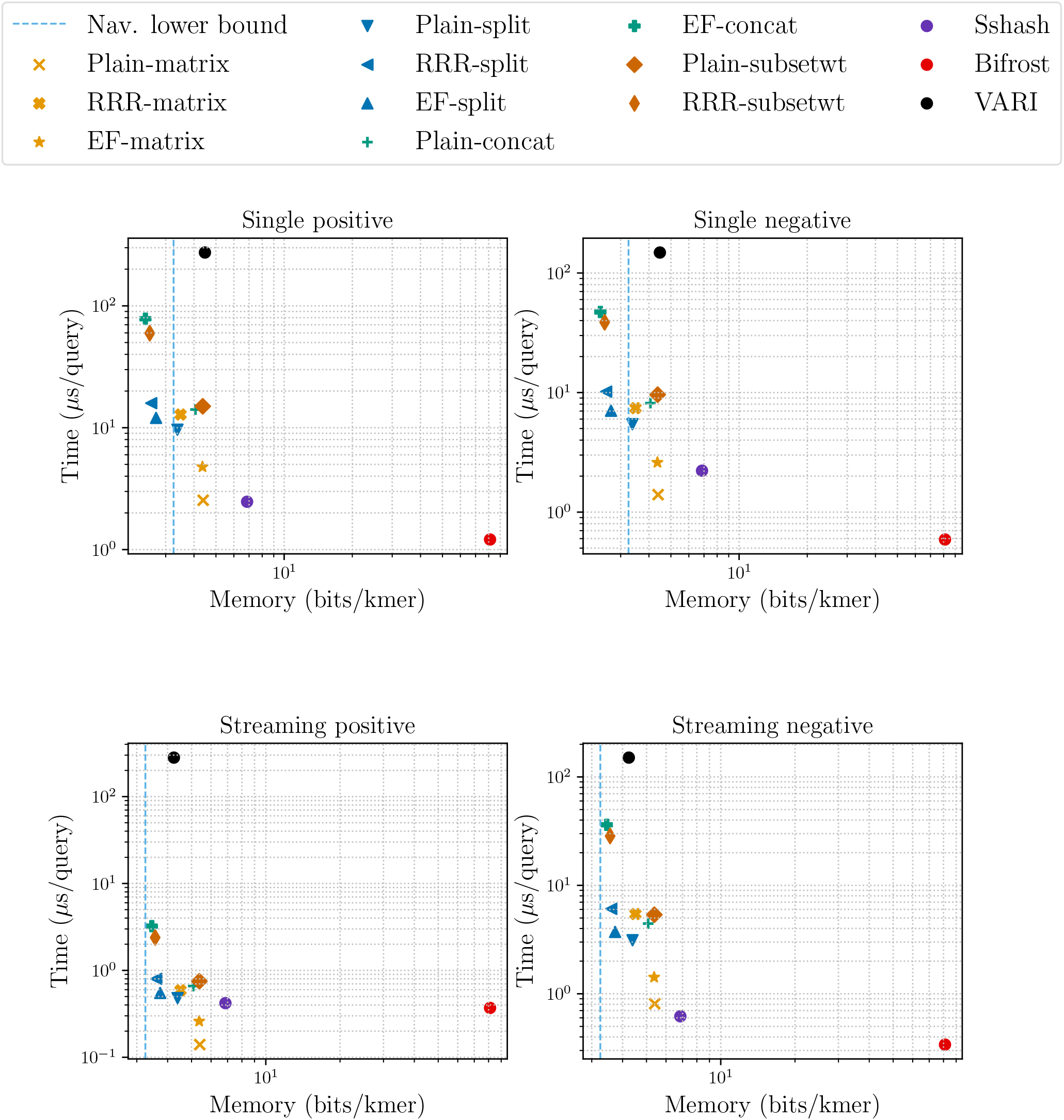
Queries on Metagenome reads.

## 7 Conclusion

While our prototypes already handsomely outperform all previous succinct de Bruijn graph implementations, we believe they can be improved in several ways, perhaps most promisingly by the replacement of wavelet trees with rank data structures specialized for small alphabet sequences. We also expect that the simplicity of the new rank-based query algorithms will make them significantly more accessible than the original BOSS, which is notoriously tricky to implement. The plain matrix, split and subset wavelet tree variants are particularly easy to implement, requiring only regular rank queries as the only non-trivial subroutine. This in contrast with the original BOSS, which also requires select and predecessor/successor queries.

## 8 Acknowledgements

We thank Giulio Pibiri for help with Sshash, Massimiliano Rossi for helpful discussion, and Harri K ä hkö nen and Daniel Cauchi for feedback on the manuscript.

## A Index construction tables

This appendix contains the tables for the results of the index construction experiments. The size is measured in bits per distinct *k*-mer (bpk) in the input data, considering reverse complements distinct.

**Table 2:** Index construction for the SARS-CoV-2 dataset.

|  | Time (s) | Mem (GB) | Size (bpk) |
| --- | --- | --- | --- |
| plain-matrix | 405 | 4.72 | 4.29 |
| rrr-matrix | 405 | 4.72 | 2.37 |
| mef-matrix | 405 | 4.72 | 3.20 |
| plain-split | 405 | 4.72 | 3.36 |
| rrr-split | 405 | 4.72 | 2.69 |
| mef-split | 406 | 4.72 | 2.78 |
| plain-concat | 406 | 4.72 | 3.90 |
| mef-concat | 406 | 4.72 | 2.04 |
| plain-subsetwt | 406 | 4.72 | 4.26 |
| rrr-subsetwt | 406 | 4.72 | 1.99 |
| sshash | 628 | 5.28 | 7.33 |
| bifrost | 3727 | 0.05 | 90.01 |
| vari | 394 | 11.23 | 4.18 |

**Table 3:** Index construction for the E. coli dataset.

|  | Time (s) | Mem (GB) | Size (bpk) |
| --- | --- | --- | --- |
| plain-matrix | 285 | 4.71 | 4.26 |
| rrr-matrix | 288 | 4.71 | 3.37 |
| mef-matrix | 293 | 4.71 | 4.07 |
| plain-split | 292 | 4.71 | 3.27 |
| rrr-split | 292 | 4.71 | 2.51 |
| mef-split | 293 | 4.71 | 2.65 |
| plain-concat | 293 | 4.71 | 3.71 |
| mef-concat | 294 | 4.71 | 2.34 |
| plain-subsetwt | 302 | 4.71 | 4.25 |
| rrr-subsetwt | 304 | 4.71 | 2.46 |
| sshash | 535 | 4.83 | 5.08 |
| bifrost | 1238 | 2.15 | 37.30 |
| vari | 310 | 11.22 | 3.18 |

**Table 4:** Index construction for the metagenomic read set.

|  | Time (s) | Mem (GB) | Size (bpk) |
| --- | --- | --- | --- |
| plain-matrix | 1552 | 8.42 | 4.30 |
| rrr-matrix | 1580 | 8.42 | 3.40 |
| mef-matrix | 1638 | 8.42 | 4.27 |
| plain-split | 1632 | 8.95 | 3.29 |
| rrr-split | 1635 | 8.75 | 2.50 |
| mef-split | 1636 | 8.79 | 2.63 |
| plain-concat | 1639 | 9.55 | 3.75 |
| mef-concat | 1653 | 9.27 | 2.34 |
| plain-subsetwt | 1747 | 12.22 | 4.28 |
| rrr-subsetwt | 1768 | 12.22 | 2.46 |
| sshash | 3808 | 8.82 | 6.85 |
| bifrost | 2194 | 29.45 | 44.10 |
| vari | 2491 | 11.20 | 4.21 |

## B Query tables

This appendix contains the tables for the results of the query experiments. The columns labeled with “+” and “-” give times for positive (found in the index) and negative queries (not found in the index) respectively. The memory is measured in bits per *k*-mer (bpk), considering a *k*-mer distinct from its reverse complement.

**Table 5:** Queries on the SARS-CoV-2 data.

|  | Single query |  |  | Streaming query |  |  |
| --- | --- | --- | --- | --- | --- | --- |
|  | + | - | mem | + | - | mem |
| | (μs) | (μs) | (bpk) | (μs) | (μs) | (bpk) |
| plain-matrix | 0.57 | 0.30 | 107.60 | 0.04 | 0.26 | 108.40 |
| rrr-matrix | 5.25 | 2.45 | 105.39 | 0.26 | 2.25 | 106.58 |
| mef-matrix | 0.59 | 0.33 | 106.19 | 0.04 | 0.26 | 107.30 |
| plain-split | 2.15 | 0.99 | 106.04 | 0.14 | 0.87 | 108.13 |
| rrr-split | 5.70 | 2.62 | 106.67 | 0.33 | 2.31 | 107.99 |
| mef-split | 2.45 | 1.14 | 106.53 | 0.14 | 0.97 | 107.67 |
| plain-concat | 2.49 | 1.14 | 107.65 | 0.14 | 0.97 | 108.66 |
| mef-concat | 28.06 | 13.50 | 106.13 | 1.37 | 12.16 | 106.71 |
| plain-subsetwt | 3.70 | 1.74 | 106.86 | 0.20 | 1.58 | 108.48 |
| rrr-subsetwt | 30.07 | 14.87 | 105.61 | 1.39 | 13.54 | 106.33 |
| sshash | 1.50 | 0.83 | 18.89 | 1.11 | 0.42 | 18.90 |
| bifrost | 0.97 | 0.48 | 141.51 | 0.95 | 0.31 | 140.93 |
| vari | 137.94 | 54.90 | 284.64 | 167.83 | 55.96 | 22.33 |

**Table 6:** Queries on the E. coli pangenome.

|  | Single query |  |  | Streaming query |  |  |
| --- | --- | --- | --- | --- | --- | --- |
|  | + | - | mem | + | - | mem |
| | (μs) | (μs) | (bpk) | (μs) | (μs) | (bpk) |
| plain-matrix | 2.13 | 0.90 | 5.25 | 0.11 | 0.55 | 6.24 |
| rrr-matrix | 10.92 | 5.23 | 4.35 | 0.50 | 4.03 | 5.35 |
| mef-matrix | 4.22 | 1.82 | 5.06 | 0.22 | 0.91 | 6.05 |
| plain-split | 7.90 | 3.66 | 4.25 | 0.36 | 1.99 | 5.25 |
| rrr-split | 13.00 | 6.41 | 3.49 | 0.61 | 4.35 | 4.49 |
| mef-split | 10.23 | 4.48 | 3.63 | 0.44 | 2.48 | 4.63 |
| plain-concat | 11.72 | 5.15 | 4.91 | 0.51 | 2.72 | 5.92 |
| mef-concat | 61.73 | 31.68 | 3.34 | 2.35 | 26.27 | 4.33 |
| plain-subsetwt | 13.00 | 6.05 | 5.23 | 0.56 | 3.28 | 6.23 |
| rrr-subsetwt | 54.71 | 28.21 | 3.44 | 2.18 | 23.16 | 4.45 |
| sshash | 1.46 | 1.45 | 5.19 | 0.23 | 0.43 | 5.19 |
| bifrost | 1.01 | 0.54 | 61.66 | 0.26 | 0.33 | 61.63 |
| vari | 216.54 | 102.70 | 5.87 | 226.22 | 103.85 | 3.38 |

**Table 7:**
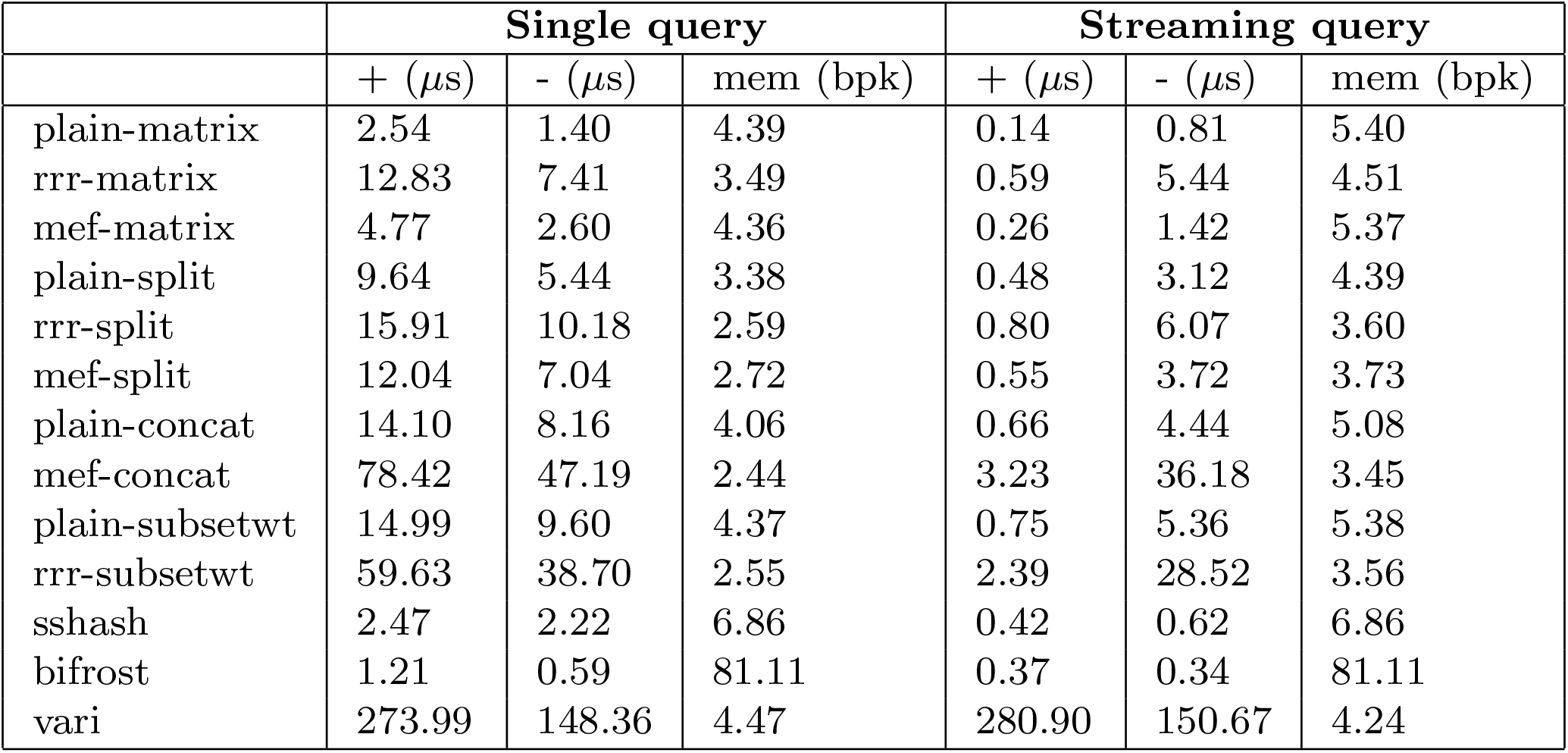
Queries on the metagenomic read set.

|  | Single query |  |  | Streaming query |  |  |
| --- | --- | --- | --- | --- | --- | --- |
| | + (μs) | - (μs) | mem (bpk) | + (μs) | - (μs) | mem (bpk) |
| plain-matrix | 2.54 | 1.40 | 4.39 | 0.14 | 0.81 | 5.40 |
| rrr-matrix | 12.83 | 7.41 | 3.49 | 0.59 | 5.44 | 4.51 |
| mef-matrix | 4.77 | 2.60 | 4.36 | 0.26 | 1.42 | 5.37 |
| plain-split | 9.64 | 5.44 | 3.38 | 0.48 | 3.12 | 4.39 |
| rrr-split | 15.91 | 10.18 | 2.59 | 0.80 | 6.07 | 3.60 |
| mef-split | 12.04 | 7.04 | 2.72 | 0.55 | 3.72 | 3.73 |
| plain-concat | 14.10 | 8.16 | 4.06 | 0.66 | 4.44 | 5.08 |
| mef-concat | 78.42 | 47.19 | 2.44 | 3.23 | 36.18 | 3.45 |
| plain-subsetwt | 14.99 | 9.60 | 4.37 | 0.75 | 5.36 | 5.38 |
| rrr-subsetwt | 59.63 | 38.70 | 2.55 | 2.39 | 28.52 | 3.56 |
| sshash | 2.47 | 2.22 | 6.86 | 0.42 | 0.62 | 6.86 |
| bifrost | 1.21 | 0.59 | 81.11 | 0.37 | 0.34 | 81.11 |
| vari | 273.99 | 148.36 | 4.47 | 280.90 | 150.67 | 4.24 |

## C Query plots

The figures in this appendix plot the data in Appendix B.

## D Additional Pseudocode

### Algorithm 3 Phase 1 of the multi-string SBWT construction algorithm.

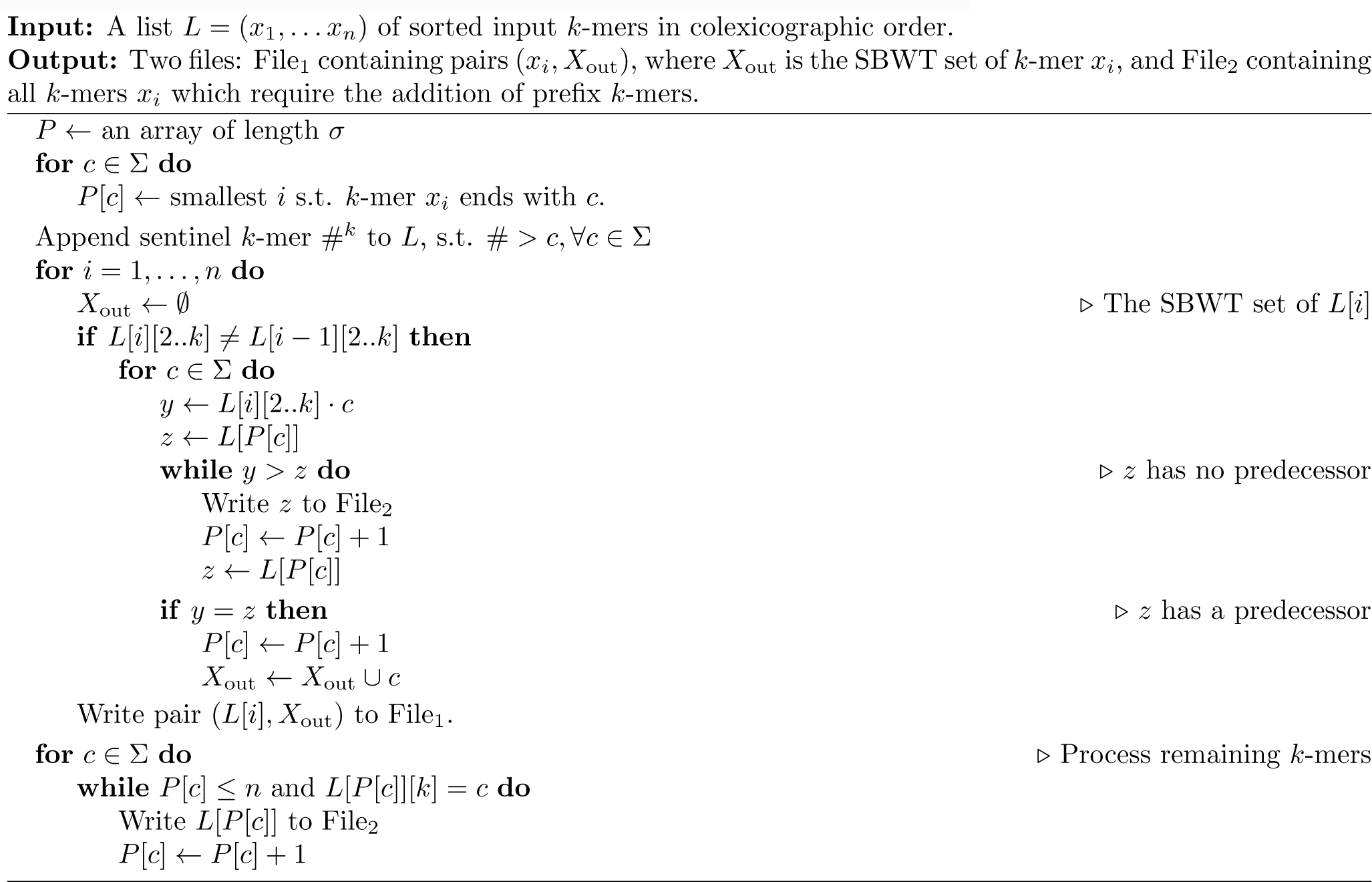

## Footnotes

1 https://github.com/cosmo-team/cosmo/tree/VARI-merge, commit 79693b7

2 https://github.com/pmelsted/bifrost/, commit 5594b5e9

3 https://github.com/jermp/sshash, commit 27852cbd

4 This is an important use case in bioinformatics, where the longer sequence typically corresponds to a so-called *read*.

## References

[1] J. Alanko, B. Alipanahi, J. Settle, C. Boucher, and T. Gagie. Buffering updates enables efficient dynamic de bruijn graphs. Computational and structural biotechnology journal, 19:4067–4078, 2021.

[2] J. Alanko, I. Slizovskiy, D. Lokshtanov, T. Gagie, N. Noyes, and C. Boucher. Syotti: Scalable bait design for dna enrichment. bioRxiv, 2021.

[3] B. Alipanahi, A. Kuhnle, S. J. Puglisi, L. Salmela, and C. Boucher. Succinct dynamic de bruijn graphs. Bioinformatics, 37(14):1946–1952, 2021.

[4] F. Almodaresi, J. Khan, S. Madaminov, M. Ferdman, R. Johnson, P. Pandey, and R. Patro. An incrementally updatable and scalable system for large-scale sequence search using the bentley-saxe transformation. Bioinformatics, 2022.

[5] C. Boucher, A. Bowe, T. Gagie, S. J. Puglisi, and K. Sadakane. Variable-order de bruijn graphs. In 2015 data compression conference, pages 383–392. IEEE, 2015.

[6] A. Bowe, T. Onodera, K. Sadakane, and T. Shibuya. Succinct de bruijn graphs. In International workshop on algorithms in bioinformatics, pages 225–235. Springer, 2012.

[7] C.-H. Chang, M.-T. Chou, Y.-C. Wu, T.-W. Hong, Y.-L. Li, C.-H. Yang, and J.-H. Hung. asBWT: memory efficient implementation of the hardware-acceleration-friendly schindler transform for the fast biological sequence mapping. Bioinformatics, 32(22):3498–3500, 2016.

[8] R. Chikhi. A tale of optimizing the space taken by de bruijn graphs. In Proc. 17th Conference on Computability in Europe (CiE), volume 12813 of LNCS, pages 120–134. Springer, 2021.

[9] R. Chikhi, A. Limasset, S. Jackman, J. T. Simpson, and P. Medvedev. On the representation of de bruijn graphs. In Proc. 18th Annual International Conference Research in Computational Molecular Biology (RECOMB), LNCS 8394, pages 35–55. Springer, 2014.

[10] R. Chikhi, A. Limasset, and P. Medvedev. Compacting de bruijn graphs from sequencing data quickly and in low memory. Bioinformatics, 32(12):i201–i208, 2016.

[11] P. E. Compeau, P. A. Pevzner, and G. Tesler. Why are de bruijn graphs useful for genome assembly? Nature biotechnology, 29(11):987, 2011.

[12] V. G. Crawford, A. Kuhnle, C. Boucher, R. Chikhi, and T. Gagie. Practical dynamic de bruijn graphs. Bioinformatics, 34(24):4189–4195, 2018.

[13] S. Deorowicz, M. Kokot, S. Grabowski, and A. Debudaj-Grabysz. Kmc 2: fast and resource-frugal k-mer counting. Bioinformatics, 31(10):1569–1576, 2015.

[14] P. Elias. Efficient storage and retrieval by content and address of static files. J. ACM, 21(2):246–260, 1974.

[15] R. Fano. On the number of bits required to implement an associative memory. Technical report, MIT, 1971.

[16] T. Gagie, G. Manzini, and J. Sirén. Wheeler graphs: A framework for bwt-based data structures. Theoretical computer science, 698:67–78, 2017.

[17] S. Gog, T. Beller, A. Moffat, and M. Petri. From theory to practice: Plug and play with succinct data structures. In International Symposium on Experimental Algorithms, pages 326–337. Springer, 2014.

[18] S. Gog and M. Petri. Optimized succinct data structures for massive data. Softw. Pract. Exp., 44(11):1287–1314, 2014.

[19] R. Grossi, A. Gupta, and J. S. Vitter. High-order entropy-compressed text indexes. In Proceedings of the fourteenth annual ACM-SIAM symposium on Discrete algorithms, pages 841–850, 2003.

[20] G. Holley and P. Melsted. Bifrost: highly parallel construction and indexing of colored and compacted de bruijn graphs. Genome biology, 21(1):1–20, 2020.

[21] Z. Iqbal, M. Caccamo, I. Turner, P. Flicek, and G. McVean. De novo assembly and genotyping of variants using colored de bruijn graphs. Nature genetics, 44(2):226–232, 2012.

[22] I. B. Jeffery, A. Das, E. O’Herlihy, S. Coughlan, K. Cisek, M. Moore, F. Bradley, T. Carty, M. Pradhan, C. Dwibedi, et al. Differences in fecal microbiomes and metabolomes of people with vs without irritable bowel syndrome and bile acid malabsorption. Gastroenterology, 158(4):1016–1028, 2020.

[23] M. Karasikov, H. Mustafa, D. Danciu, M. Zimmermann, C. Barber, G. Rätsch, and A. Kahles. Metagraph: Indexing and analysing nucleotide archives at petabase-scale. BioRxiv, 2020.

[24] J. Kärkkäinen, D. Kempa, and S. J. Puglisi. Hybrid compression of bitvectors for the FM-index. In Proc. DCC, pages 302–311. IEEE, 2014.

[25] J. Khan, M. Kokot, S. Deorowicz, and R. Patro. Scalable, ultra-fast, and low-memory construction of compacted de bruijn graphs with cuttlefish 2. bioRxiv, 2021.

[26] J. Khan and R. Patro. Cuttlefish: fast, parallel and low-memory compaction of de bruijn graphs from large-scale genome collections. Bioinformatics, 37(Supplement 1):i177–i186, 2021.

[27] M. Kokot, M. Dlugosz, and S. Deorowicz. Kmc 3: counting and manipulating k-mer statistics. Bioinformatics, 33(17):2759–2761, 2017.

[28] D. Ma, S. J. Puglisi, R. Raman, and B. Zhukova. On Elias-Fano for rank queries in FM-indexes. In 31st Data Compression Conference (DCC), pages 223–232. IEEE, 2021.

[29] N. Maillet, C. Lemaitre, R. Chikhi, D. Lavenier, and P. Peterlongo. Compareads: comparing huge metagenomic experiments. In BMC bioinformatics, volume 13, pages 1–10. Springer, 2012.

[30] T. Mäklin, T. Kallonen, J. Alanko, Ø. Samuelsen, K. Hegstad, V. Mäkinen, J. Corander, E. Heinz, and A. Honkela. Bacterial genomic epidemiology with mixed samples. Microbial genomics, 7(11), 2021.

[31] C. Marchet, C. Boucher, S. J. Puglisi, P. Medvedev, M. Salson, and R. Chikhi. Data structures based on k-mers for querying large collections of sequencing data sets. Genome Research, 31(1):1–12, 2021.

[32] C. Marchet, M. Kerbiriou, and A. Limasset. BLight: efficient exact associative structure for k-mers. Bioinformatics, 37(18):2858–2865, 04 2021.

[33] M. D. Muggli, A. Bowe, N. R. Noyes, P. S. Morley, K. E. Belk, R. Raymond, T. Gagie, S. J. Puglisi, and C. Boucher. Succinct colored de Bruijn graphs. Bioinformatics, 33(20):3181–3187, Oct. 2017.

[34] J. I. Munro. Tables. In Proc. 16th Conference on Foundations of Software Technology and Theoretical Computer Science, LNCS 1180, pages 37–42. Springer, 1996.

[35] G. Navarro and E. Providel. Fast, small, simple rank/select on bitmaps. In R. Klasing, editor, Experimental Algorithms - 11th International Symposium, SEA 2012, Bordeaux, France, June 7-9, 2012. Proceedings, volume 7276 of Lecture Notes in Computer Science, pages 295–306. Springer, 2012.

[36] D. Okanohara and K. Sadakane. Practical entropy-compressed rank/select dictionary. In Proc. Ninth Workshop on Algorithm Engineering and Experiments (ALENEX). SIAM, 2007.

[37] B. D. Ondov, T. J. Treangen, P. Melsted, A. B. Mallonee, N. H. Bergman, S. Koren, and A. M. Phillippy. Mash: fast genome and metagenome distance estimation using minhash. Genome biology, 17(1):1–14, 2016.

[38] J. Pell, A. Hintze, R. Canino-Koning, A. Howe, J. M. Tiedje, and C. T. Brown. Scaling metagenome sequence assembly with probabilistic de bruijn graphs. Proceedings of the National Academy of Sciences, 109(33):13272–13277, 2012.

[39] G. E. Pibiri. Sparse and skew hashing of k-mers. bioRxiv, 2022.

[40] A. Rahman and P. Medvedev. Representation of k-mer sets using spectrum-preserving string sets. In International Conference on Research in Computational Molecular Biology, pages 152–168. Springer, 2020.

[41] R. Raman, V. Raman, and S. R. Satti. Succinct indexable dictionaries with applications to encoding k -ary trees, prefix sums and multisets. ACM Transactions on Algorithms, 3(4):43, Nov. 2007.

[42] E. A. Rødland. Compact representation of k-mer de bruijn graphs for genome read assembly. BMC bioinformatics, 14(1):1–19, 2013.

[43] S. Vigna. Quasi-succinct indices. In Proc. Sixth ACM International Conference on Web Search and Data Mining (WSDM), pages 83–92. ACM, 2013.

